# A key residue of the extracellular gate provides quality control contributing to ABCG substrate specificity

**DOI:** 10.1101/2024.04.22.590559

**Authors:** Jian Xia, Alexandra Siffert, Odalys Torres, Joanna Banasiak, Konrad Pakuła, Jörg Ziegler, Sabine Rosahl, Noel Ferro, Michał Jasiński, Tamás Hegedűs, Markus M. Geisler

## Abstract

For G-type ATP-binding cassette (ABC) transporters, a hydrophobic “di-leucine motif” as part of a hydrophobic extracellular gate has been described to separate a large substrate-binding cavity from a smaller upper cavity and proposed to act as a valve controlling drug extrusion.

Here, we show that an L704F mutation in the hydrophobic extracellular gate of Arabidopsis ABCG36/PDR8/PEN3 uncouples the export of the auxin precursor indole-3-butyric acid (IBA) from that of the defense compound camalexin (CLX). Molecular dynamics simulations reveal an increase in free energy and pulling forces for CLX at both the entrance and exit sites of ABCG36^L704F^, respectively, providing a mechanistic rationale for the transport discrimination of CLX. Mutagenesis of L704 to tyrosine allows export of structurally related non-ABCG36 substrates, indole-3-acetic acid (IAA) and indole, suggesting an allosteric communication between the extracellular gate and the central substrate binding pocket.

In summary, our work supports the conclusion that L704 is a key residue of the extracellular gate that provides a final quality control contributing to ABCG substrate specificity.

## Introduction

ABC proteins represent an ancient and versatile transport system operating in all living organisms ^1–3^. Most ABC transporters are primary pumps, which utilize the energy of ATP-dependent hydrolysis to transport various substrates across cellular membranes ^4^. Ectopic or dysregulated over-expression of certain ABC transporters often contributes to multidrug resistance (MDR) phenomena due to non-selective translocation of various drugs and thus can be a severe complication for treating cancer as well as for microbial and parasitic infections ^5^. Three human ABC transporters, ABCB1 (Multidrug resistance protein 1, MDR1; P-glycoprotein 1, Pgp1), ABCC1 (Multidrug resistance-associated protein 1, MRP1) and ABCG2 (Breast cancer resistance protein, BCRP), are thought to commonly promote or cause MDR in a variety of therapeutic cancer settings.

In higher plants, the ABC family is expanded twice to four times compared to other organisms, including mammals or their ancestral microalgae ^6,7^. This has led to the concept that ABC transporters multiplied during evolution and acquired novel functions that allowed plants to adapt to a sessile and terrestrial lifestyle ^8^. This idea is indirectly supported by the finding that many plant ABC transporters transport growth hormones, like auxins and cytokinins playing a crucial role in post embryonic development ^7^. Up to date, research on plant ABCs has focused on ABCBs because a subset was shown to catalyze the transport of auxin and to be determinants of plant development ^9–12^. Recently, the focus has shifted toward PLEIOTROPIC DRUG RESISTANCE (PDR)-type ABC transporters, which are full-size ABC transporters of the G-type and found mostly in plants and fungi ^6,13^. One of the most investigated plant PDR to date is ABCG36/PENETRATION3 (PEN3)/PDR8, for which a range of structurally unrelated, putative substrates were identified ^13,14^. Besides the auxin precursor, indole-3-butyric acid (IBA; ^13^), ABCG36 was shown to export two indolic compounds functioning in defense signaling ^15^. Moreover, ABCG36 was characterized as an exporter of the antimicrobial, indolic compound camalexin (CLX; ^14^). Based on the physiological importance of these substrates (IBA and CLX) for Arabidopsis, it was suggested that ABCG36 functions at the cross-road between plant growth and pathogen defense, which represents one of the significant tradeoffs that plants have to balance ^16^. The role of ABCG36 in these two programs was recently shown to be regulated via protein phosphorylation by the leucine-rich repeat receptor kinase QIAN SHOU KINASE1. ABCG36 phosphorylation unilaterally represses IBA export, promoting CLX export by ABCG36 conferring pathogen resistance ^14^.

Significant resources were dedicated to uncovering the structure of ABCG proteins to understand their function and substrate recognition. The first high-resolution ABCG crystal structures of the human half-size transporter heterodimer, ABCG5/ABCG8 ^17^, and cryo-EM structures of the ABCG2 homodimer ^18,19^ revealed that despite an overall similar transport mechanism, ABCG transporters embody a transmembrane domain structure later described as type V ^20^, that is significantly different compared to type IV transporters, represented by ABCB, ABCC and ABCD subfamily members. In the core of the substrate promiscuous ABCG2 homodimer, a hydrophobic “di-leucine motif” was described to separate a larger lower intracellular cavity - serving as a binding region for substrates and inhibitors - from a smaller upper cavity ^18^. Later, this hydrophobic “leucine plug” was suggested to act as a valve to control drug extrusion ^21^. Furthermore, the ABCG2 structure contains a polar “extracellular roof” with a compact semi-closed architecture, which supposedly acts as a barrier for substrate release from the upper cavity ^21^. In contrast, ABCG5/G8 exhibits a collapsed cavity of insufficient size to hold the substrates ^22^ and leucine residues of the putative valve are substituted by hydrophobic residues phenylalanine and methionine, respectively. The functional implications of these residue differences are unclear but the larger aromatic phenylalanine side chain was speculated to be critical for sterol selectivity ^22,23^.

The structure of the yeast pleiotropic drug resistance transporter PDR5 was resolved by single-particle cryo-EM ^24^. In contrast to other ABCGs, one-half of the transporter remains nearly invariant throughout the cycle, and the other half undergoes changes transmitted across inter-domain interfaces to support the transport process by peristaltic motions. In an AlphaFold2-derived structure of PDR-type ABCG46 from the model crop plant *Medicago truncatula*, an unusually narrow transient access path to the central cavity constitutes an initial filter responsible for the selective translocation of phenylpropanoids ^25^. Mutation of F562 located in TH2 (Transmembrane Helix 2) affected the viability of the transient access path through helix rearrangements, profoundly affecting the selectivity of MtABCG46.

Recently, three independent studies published cryo-EM structures of the Arabidopsis half-size transporter, ABCG25, working as a homodimeric exporter of the plant hormone abscisic acid (ABA) ^26–28^. The interface between ABA and ABCG25 is mainly formed by hydrophobic interactions with the residues of TH2 and TH5a ^27^. Interestingly, here ABA translocation seems to be controlled by two gates: a “cytoplasmic gate” formed by residue F453 from TH2/TH2′, and an “apoplastic gate” formed by Y565 from TH5a/TH5a′ ^27^. The latter is a functional homolog of the hydrophobic “leucin plug” described for human ABCG2 ^18,21^.^1^

Based on pathogen susceptibility and IBA sensitivity assays, a specific point mutation in *ABCG36/PEN3* of Arabidopsis was described to uncouple ABCG36 functions in IBA-stimulated root growth, callose deposition and pathogen-inducible salicylic acid accumulation from ABCG36 activity in pathogen defense. It was suggested that this uncoupling in the *abcg36-5*/*pen3-5* allele might be caused by lack of export of major relevant defense compounds (such as CLX), while IBA export was thought to be preserved ^29^. Interestingly, the *abcg36-5* allele encodes for a L704F change in elongation of TH5 (misannotated as TH4 in Lu et al. (2015)) of ABCG36.

In this study, we uncover that uncoupling of ABCG36-mediated function in IBA-stimulated root growth from its transport role in defense in the *abcg36-5* allele (L704F) is due to unilateral loss of CLX transport capacity, while IBA transport is preserved. L704 (together with F1375) is a *Brassicaceae* family-specific key residue of an extracellular, hydrophobic gate that provides quality control of substrate specificity in ABCG36.

## Results

### L704F mutation in ABCG36 uncouples export of IBA and camalexin

Recent work has established the engagement of multiple ABCG36 substrates in different ABCG36-dependent critical, biological processes, including growth and defense ^13,14,29–36^. The *abcg36-5* allele uncouples ABCG36 functions in IBA-stimulated root growth from ABCG36 activity in extracellular defense ^29^ but the underlying molecular determinants are unclear. This unique IBA-uncoupling of the *abcg36-5* allele was originally found by using an IBA-mediated primary root growth inhibition phenotype in the presence of toxic µM concentrations ^14,29,37^.

In order to verify these findings, we replicated the originally described detoxification assay ^37^ under our conditions, and included the *abcg36-4* and *abcg36-6* loss-of-function alleles as negative controls. The first is a T-DNA insertional null allele, while the second carries an A1357V substitution in TH11 ^29^ (Suppl. Fig. 3e). As shown before ^29^, *abcg36-4* and *abcg36-6* alleles but not *abcg36-5* exhibited reduced root lengths on IBA in comparison with their Wt backgrounds (Suppl. Fig. 1a-b). A repetition of this assay on cytotoxic concentrations of CLX ^14^, indicated as expected a hypersensitivity for *abcg36-4*, *abcg36-6* and *abcg36-5* alleles in comparison to their corresponding Wt backgrounds, Col-0 and Gl1, respectively (Suppl. Fig. 1c-d).

These results suggested that selective uncoupling of IBA detoxification was most likely due to the lack of CLX export capacities on *abcg36-5* ^13^. In order to test this possibility, we firstly quantified IBA and CLX export from leaf mesophyll protoplasts prepared from the leaves of Wt and mutant plants after loading of ^3^H-IBA or ^3^H-CLX, respectively ^12,14^. Results indicated significantly reduced CLX but not IBA export from *abcg36-5* cells, while both *abcg36-4* and *abcg36-6* showed significantly reduced IBA and CLX export (Fig. 1a-b). For the chemically related auxin, IAA, previously reported not to be transported by ABCG36 ^14,38^, as well as the diffusion control benzoic acid (BA) and indole, no significant differences to Wt were found (Fig. 1c; Suppl. Fig. 2a-b).

**Figure 1.**
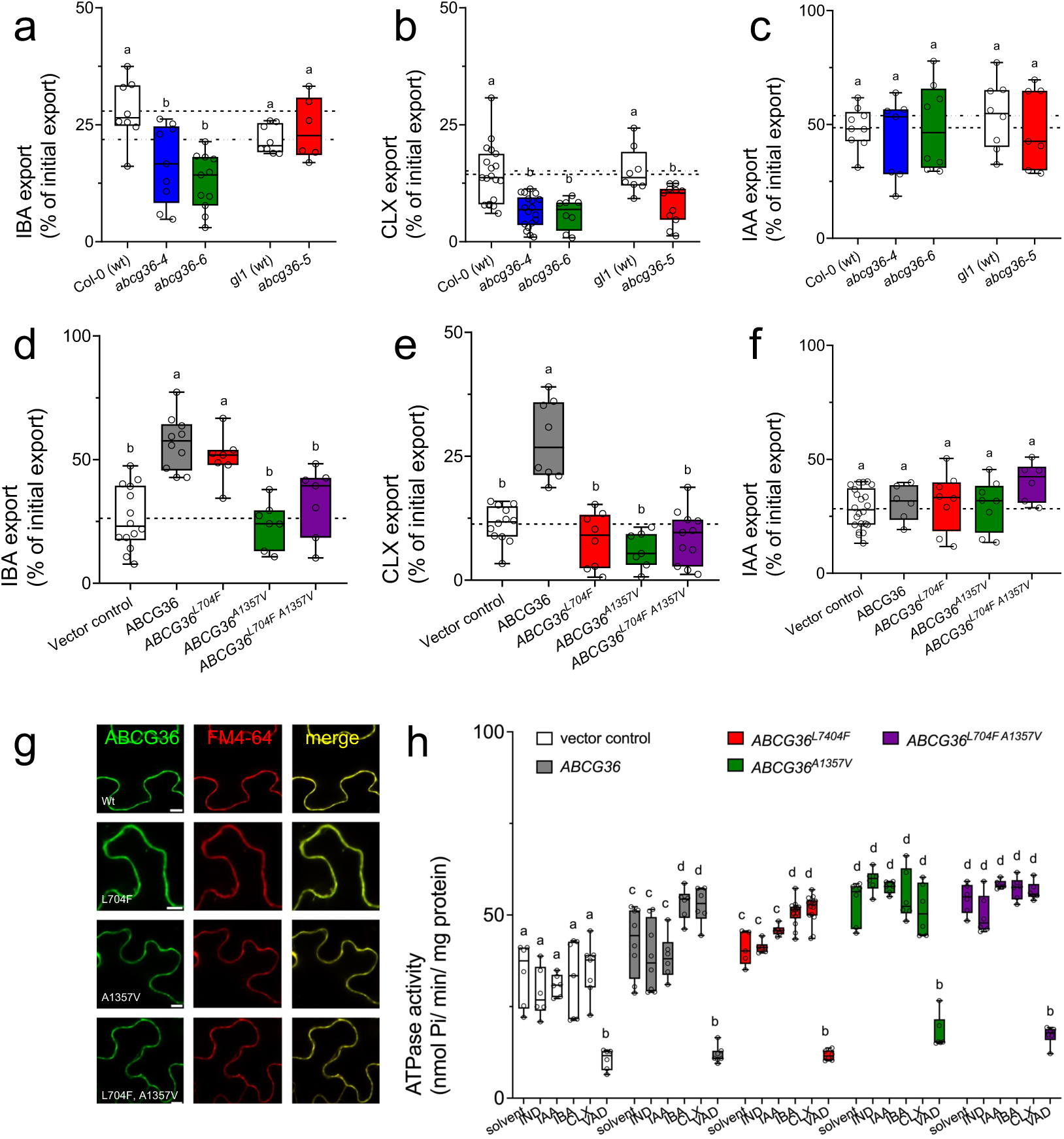
L704F mutation in ABCG36 uncouples IBA from camalexin export. **a-c.** IBA (**a**) CLX (**b**) and IAA (**c)** export from *Arabidopsis* protoplasts prepared from indicated *ABCG36* loss-of-function alleles. Significant differences (*p* < 0.05) of means ± SE (n ≥ 7 independent protoplast preparations) were determined using Brown-Forsythe and Welch ANOVA and are indicated by different lowercase letters. **d-f.** IBA (**d**) CLX (**e**) and IAA (**f)** export from *N. benthamiana* protoplasts after transfection with indicated mutant versions *ABCG36*. Significant differences (*p* < 0.05) of means ± SE (n ≥ 7 independent protoplast preparations) were determined using Brown-Forsythe and Welch ANOVA and are indicated by different lowercase letters. **g.** Confocal imaging of GFP-tagged versions of mutated *ABCG36* after transfection of *N. benthamiana* leaves. Short treatment of FM4-64 was used as PM markers; bar, 50 μm. **h.** ATPase activity of microsomal fractions prepared from tobacco leaves transfected with vector control or indicated mutant versions of *ABCG36* measured at pH 9.0 in the presence of 50 μM indole, IAA, IBA, CLX, or ortho-vanadate. Significant differences (*p* < 0.05) of means ± SE (n = 3 independent transfections and microsomal preparations) were determined using Two-way ANOVA and are indicated by different lowercase letters.

To secondly verify these Arabidopsis data in a heterologous system allowing to exclude off-target effects by mutational events, we functionally expressed Wt and mutant versions, which contained the exact point mutations in *ABCG36-GFP* as in the described *abcg36* alleles, in tobacco by using agrobacterium-mediated leaf transfection ^39^. As expected, the L704F (*abcg36-5*) variant had no effect on ABCG36-mediated IBA export but abolished ABCG36-mediated CLX export to vector control level (Fig. 1d-e). Further, the A1357V (*abcg36-6*) allele was not able to transport neither of the substrates (Fig. 1d-e). Finally, combining both substitutions had no significant additional impact excluding additive effects (Fig. 1d-e). As before in Arabidopsis, mutations had no significant effect on the export of IAA, BA, and indole as being non-ABCG36 substrates (Fig. 1f, Suppl. Fig. 2c-d) or on PM presence or PM expression level in tobacco based on confocal imaging (Fig. 1g).

We also tested the effect of these mutations on ABCG36 ATPase activities measured on isolated microsomes prepared from transfected tobacco at pH 9, where ATPase activity of H^+^-ATPases is neglectable, excluding indirect effects^14^. Recently, ABCG36 ATPase activity was found to be stimulated by its substrates, IBA and CLX, but not by non-transported indolic compounds, IAA, or indole ^14^. ATPase activity of ABCG36^L704F^ (Fig. 1h) was comparable to Wt and showed likewise a stimulation by IBA and CLX. Activities of ABCG36^A1357V^ were slightly enhanced but lacked substrate stimulation. The latter is of interest because recently L554A substitution of the homologous residue in human ABCG2 resulted in an unstable protein, while L555A exchange showed lower substrate-stimulated ATPase activity, but twice the translocation activity of the wild-type (Manolaridis et al., 2018).

Thirdly, in order to exclude ecotype-specific (Col-0 vs. Gl1 Wt) differences in root elongation and/or transport, we complemented the T-DNA insertion loss-of-function allele *abcg36-4* with single and double point-mutated, genomic versions of *ABCG36* (*ABCG36:ABCG36-GFP*) expressed under its native promoter ^36^. Like Wt ^14^, two independent lines of *ABCG36^L704F^*fully complemented the hypersensitivity of *abcg36-4* roots toward IBA and its incompetence of IBA export (Fig. 2a, c), while this was not found for CLX (Fig. 2b, d). As seen with chemically generated point mutations (Fig. 2a-d; ^29^), *ABCG36^A1357V^* and *ABCG36^L704F, A1357V^*lines did not complement the *abcg36-4* mutant. As before, these mutations did not significantly alter PM expression and polarity of ABCG36-GFP in Arabidopsis (Fig. 2e).

**Figure 2.**
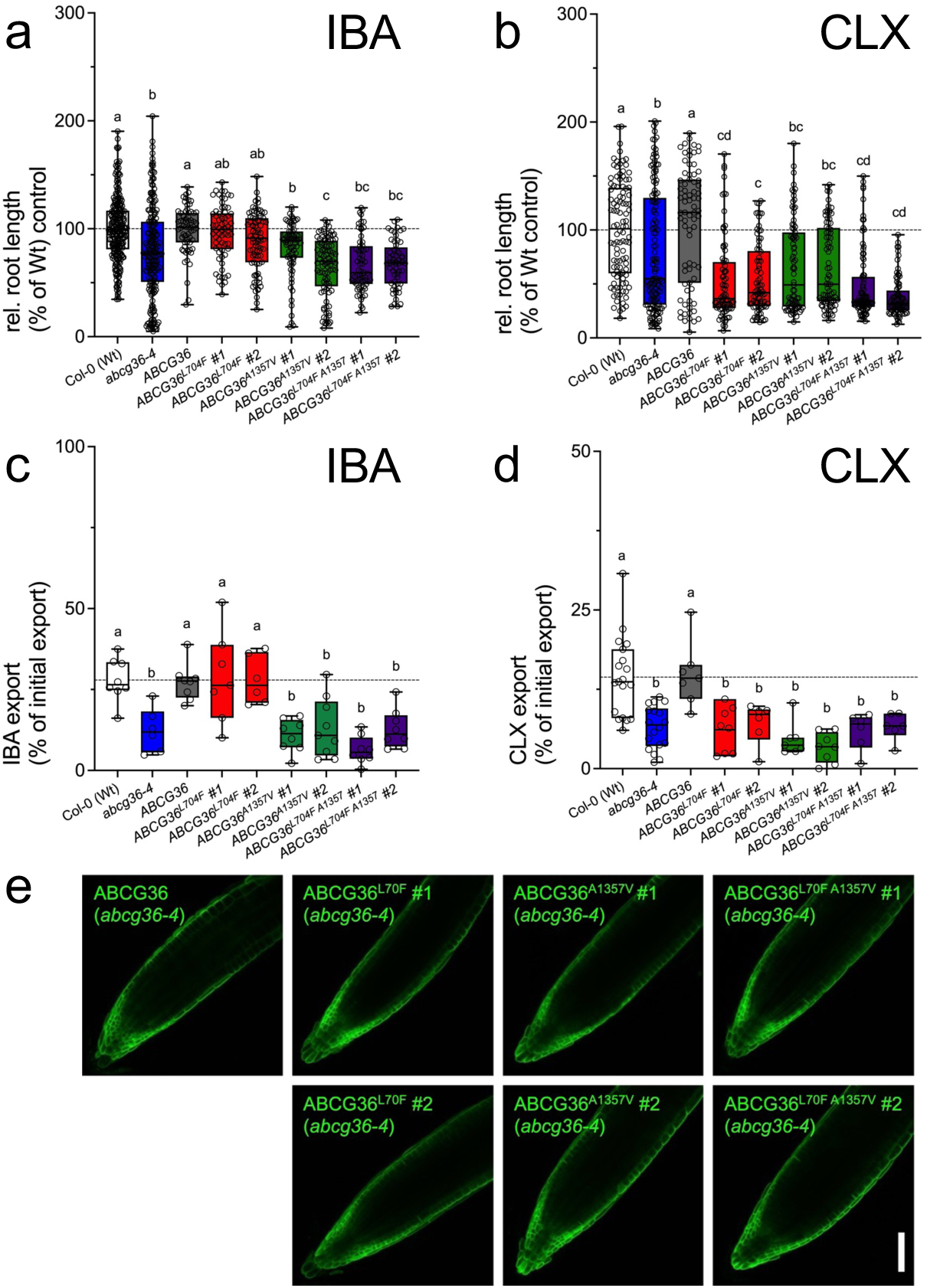
Uncoupling of IBA from camalexin export is ecotype-independent. **a-b.** Relative root length of two independent Arabidopsis *abcg36-4* lines complemented with indicated mutant versions of *ABCG36* grown for 12 days on 7.5 μM IBA (**a**) or 5 ug/ml CLX (**b**); Wt growth is set to 100%. Significant differences (*p* < 0.05) of means ± SE (n = 3 independent experiments with each 10-16 seedlings) were determined using Ordinary one-way ANOVA and are indicated by different lowercase letters. **c-d.** IBA (**c**) and CLX (**b**) export from two independent Arabidopsis *abcg36-4* lines complemented with indicated mutant versions of *ABCG36*. Significant differences (*p* < 0.05) of means ± SE (n ≥ 7 independent protoplast preparations) were determined using Brown-Forsythe and Welch ANOVA and are indicated by different lowercase letters. **e.** Confocal imaging of Arabidopsis *abcg36-4* lines complemented with indicated mutant versions of *ABCG36*; bar, 50 μm.

These datasets clearly demonstrate that uncoupling of ABCG36-mediated function in IBA-stimulated root growth from its transport role in defense in the *abcg36-5* allele (L704F) is due to unilateral loss of CLX transport capacity, while IBA transport is preserved.

### ABCG36 might own a signaling function during infection

For *abcg36-5* and *abcg36-6* alleles, mixed sensitivities toward shoot pathogens have been reported: while both alleles retain susceptibility to the host-adapted fungus, *Golovinomyces orontii*, they are fully defective in defense to non-adapted powdery mildews ^29^. In order to test how these point mutations perform upon root infection, we treated them with *Fusarium oxysporum*, a well-characterized, root vascular fungal pathogen that causes wilt disease in several plant species, including *Arabidopsis thaliana* ^40^. This pathosystem was recently used to uncover a phospho-switch balancing IBA and CLX transport capacities ^14^. In line with reduced CLX export capacities caused by L704F and A1357V mutations (Figs. 1-2), both alleles were like KO mutants ^14^ hyper-susceptible toward *Fusarium* infection based on quantification of leaf symptoms (Fig. 3a-b).

**Figure 3:**
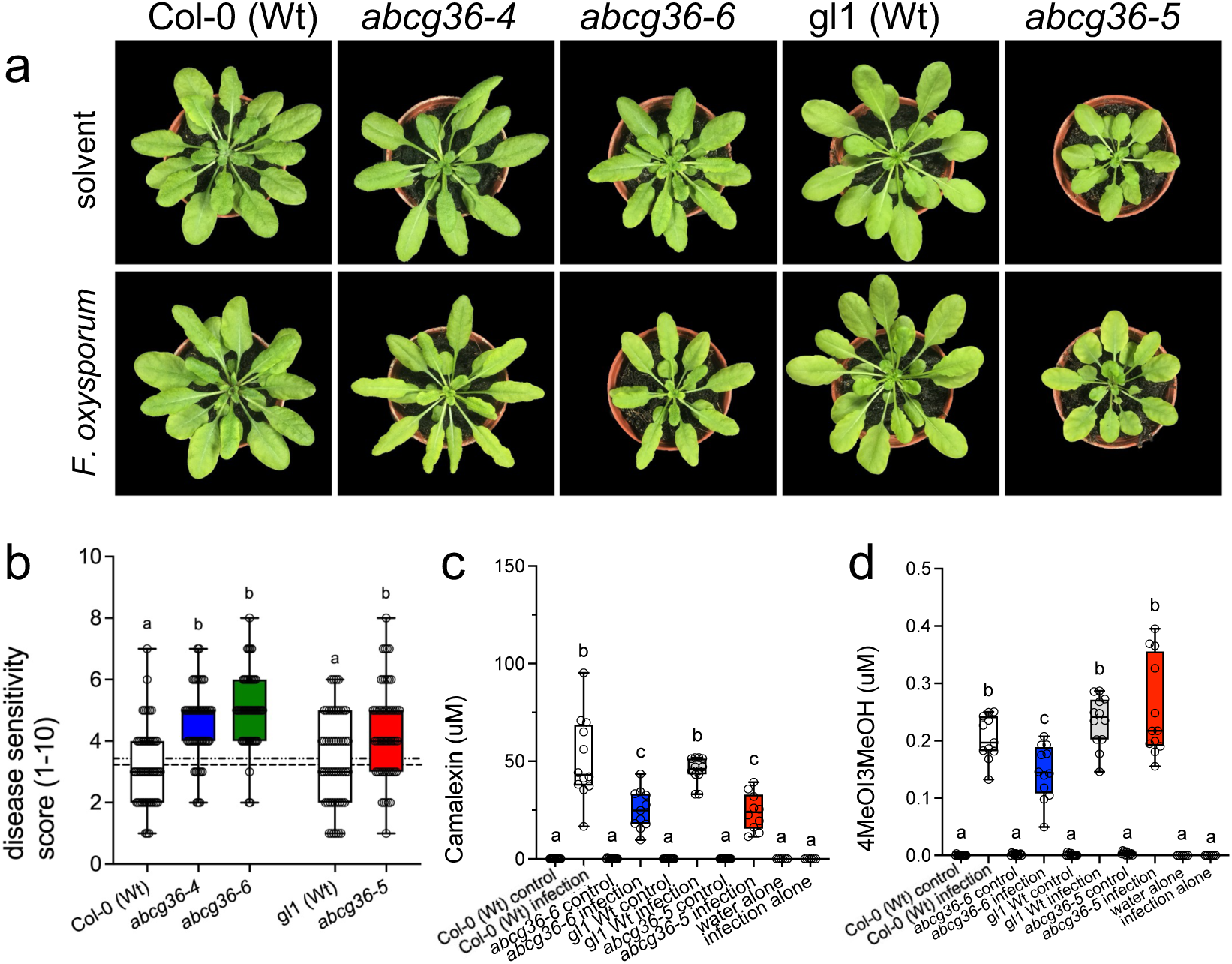
ABCG36 has a sensor-like function during infection. **a-b.** 5-week-old plants grown on soil were watered with buffer (untreated control) or with Fo (Fo699; 10^7^ conidia/ml). Representative plants are pictured (**a**) and disease symptoms were quantified (**b**) using a scale from 0-10 ^14^. Significant differences (*p* < 0.05) of means ± SE (n = 3 independent infection series with each 15-20 single plants) were determined using Brown-Forsythe and Welch ANOVA with Dunnet’s T3 multiple comparison test and are indicated by different lowercase letters. **c-d.** Extracellular levels of camalexin (**c**) and 4MeOI3M (**d**) on *P. infestans*-inoculated *Arabidopsis* leaves determined by non-targeted UPLC-ESI-QTOF-MS. Significant differences (*p* < 0.05) of means ± SE (n ≥ 7 with each 30 droplets) were determined using Ordinary one-way ANOVA and are indicated by different lowercase letters.

In order to be able to quantify exported ABCG36 substrates upon infection in these alleles, we changed back to a leaf pathosystem using established drop-inoculation of leaves with the oomycete *Phytophthora infestans* ^14,30^. As described previously, water controls of both Wt ecotypes showed very low surface levels of CLX ^14,30^ that were not significantly different from *abcg36* alleles (Fig. 3c). However, surface drops that contained *P. infestans* zoospores revealed highly enhanced CLX levels in Wt compared to the non-infected control, while CLX concentrations collected from *abcg36-5* and *abcg36-6* leaves were strongly reduced in comparison to the infected Wts, respectively (Fig. 3c).

Because it is very difficult to reliably detect IBA in the infection droplets, most likely due to its low abundance ^41^, we looked at the ABCG36 substrate, 4-methoxyindol-3-yl-methanol (4MeOI3M), which based on competition experiments is transported in an IBA-specific manner ^14,30^. In agreement with IBA transport experiments (Figs. 1-2), 4MeOI3M export upon infection was significantly reduced in the transport-dead *abcg36-6* allele but not in *abcg36-5* (Fig. 3d) shown to retain IBA export (Figs. 1-2) and IBA-mediated root elongation ^29^.

Reduced CLX and 4MeOI3M export for *abcg36-6* but only a reduction of CLX export for *abcg36-5* in comparison to the corresponding Wt seems to be intuitively correct but is on a first view in conflict with previously published data showing strongly enhanced CLX and reduced 4MeOI3M presence, respectively, in infection droplets for both *abcg36-3* and *abcg36-4* T-DNA insertional, loss-of-function mutants upon infection ^14^. A rational for this behavior was provided by the finding that ABCG40, a putative CLX and a *bona fide* IBA exporter ^31^, was strongly upregulated on the protein level in the *abcg36-3* allele upon infection ^14^. A reasonable and attractive explanation for this contradiction can be found in the concept that this over-compensation by ABCG40 (and eventually other related isoforms; ^14^) might not take place in mutated, PM expressed versions of ABCG36 (as found for *abcg36-5* and *abcg36-6* alleles; Fig. 3c) but only in the absence of the ABCG36 protein as proven for *abcg36-3* and *abcg36-4* null alleles ^29^.

In summary, this dataset supports on one hand the unilateral absence of CLX transport for the *abcg36-5* allele in a root-based pathosystem but on the other indicates a previously unknown signaling function for ABCG36 in an interplay with closely related ABCG isoforms.

### The L704 mutation is part of the extracellular gate and influences the transport process primarily by allosteric effects

To provide a mechanistic explanation for the ability of ABCG36 to discriminate IBA and IAA and for the unilateral substrate discrimination between IBA and CLX found for the L704F variant, we employed various *in silico* methods. Since *in silico* docking of substrates to ABCG2 suggested several drug-binding spots along the substrate translocation pathway ^42^, we performed metadynamics simulations to study the translocation process. However, because such a large protein system likely prohibits reaching full convergence, we supplemented these calculations with pulling simulations.

The inside-closed AlphaFold2 structure of ABCG36 was used in all of our simulations since this structure is the closest to the transport-competent conformation (Suppl. Fig. 3 ^43^). Interestingly, sequence analysis and mapping of the *abcg36-5* mutation on this ABCG36 structure confirmed that L704 is part of the “leucine valve” terminating the substrate funnel, which is formed by extensions of transmembrane helices TH2, TH5, TH8, and TH11 (Figs. 4c and 5a; Suppl. Fig. 7 ^21^). F703/F1374 and L704/F1375 of ABCG36 correspond to L554 and L555 of human ABCG2, respectively (Suppl. Fig. 7a) and protrude from TH5 and TH11, respectively (Suppl. Fig. 7). A1357, the cause of the *abcg36-6* mutation, is also part of TH11 but is located towards the substrate entry side (Suppl. Fig. 3e).

**Figure 4:**
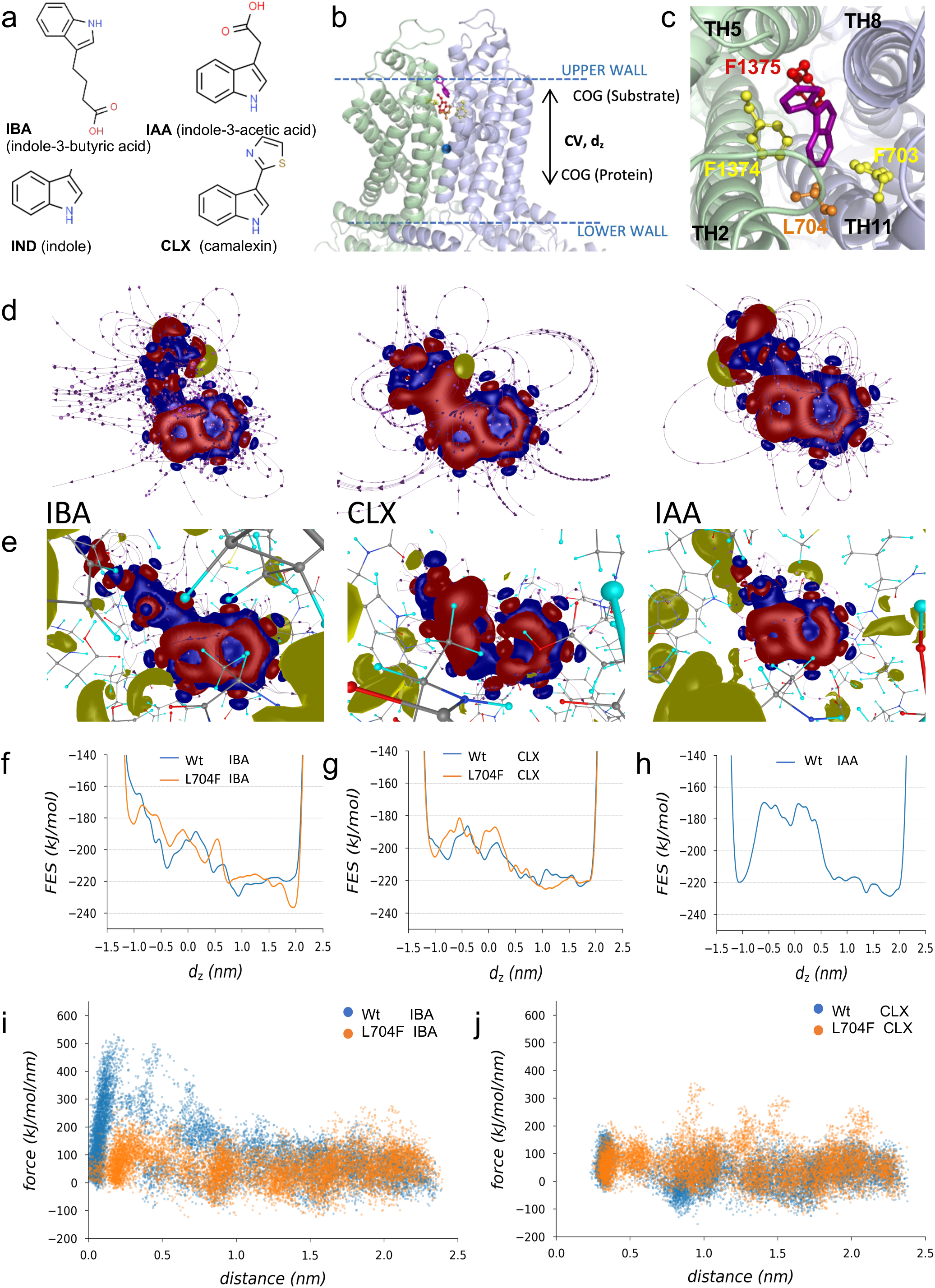
L704 is part of the extracellular gate of ABCG36 and L704F increases free energy surface for camalexin in the entrance region. **a.** Chemical formulas for potential ABCG36 substrates employed in this study. **b.** The reaction coordinate (collective variable, CV) for metadynamics is demonstrated using the side view of the ABCG36 structure. The z component (orthogonal to the membrane bilayer) of the distance between the center of geometry (COG; a small, blue circle, defined by the TH regions of TH2, TH5, TH8, and TH11) and the substrate defines the value of CV, limited by an upper and a lower wall to prohibit the escape of the small molecules from the translocation pathway. **c.** THs of ABCG36 are shown from the extracellular space. Extracellular gate residues are highlighted by sticks and balls. Docked IBA is shown using stick representation. **d-e.** Electronic structural deformation, the electric field lines of IBA, CLX and IAA before (**d**) and after binding to the ABCG36 substrate pocket (**e**). The electronic binding energies and the results displayed here were obtained by *ab initio* calculations using DFT methods. **f-h**. Free energy surfaces (FES) were calculated from metadynamics simulations with complexes of IBA (f), CLX (g), and IAA (h) with Wt and L704F ABCG36, respectively. **i-j**. Pulling simulations of IBA (i) and CLX (j) from the central binding pocket to the extracellular space were performed in simulations with Wt and L704F ABCG36, respectively. Force distributions are plotted.

**Figure 5:**
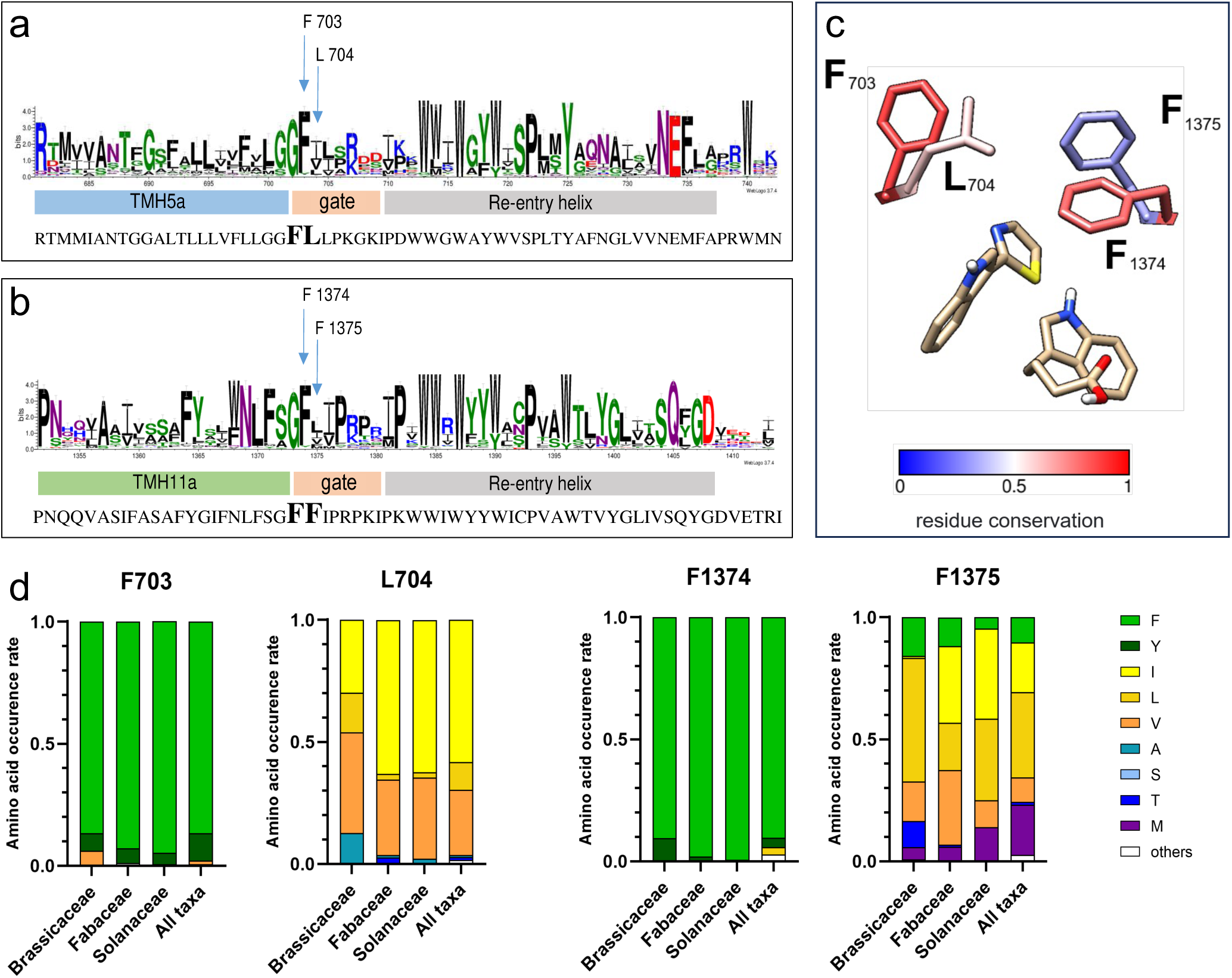
Evolutionary analysis of key residues in the putative extracellular gate. **a-b.** WebLogo presentation of the extracellular gate regions of TH5 (a) and TH11 (b) of 1853 full-size ABCGs. The x-axis displays the position of amino acids in the multiple sequence alignments according to AtABCG36. **c.** L704 and F1375 are less conserved than F703 and F1374 in full-size ABCG sequences, the positions of ABCG substrates, IBA, and CLX are indicated. **d.** Occurrence of amino acids corresponding to F703, L704, F1374, and F1375 in Arabidopsis ABCG36 in full-size ABCG sequences of indicated taxa.

*Ab initio* calculations using density functional theory (DFT) allowed to calculate electronic structural deformation and the electric field lines of IBA, CLX and IAA before and after binding to the ABCG36 substrate pocket (Fig. 4d-e). For IBA and CLX, the electronic structure remained stable though revealing changes in field lines. In contrast, the binding of IAA to the ABCG36 pocket leads to significant changes in the electronic structure of IAA, particularly in its side chain (Fig. 4d-e). This is accompanied by a partitioning of the electronic deformation within the interatomic bonding structure, resulting in a decrease in internal molecular stability. The buffer region, which corresponds to the coupling between the side chain and the IAA ring system, also exhibits the same phenomenon. The electronic structure of the internal bonds is not retained after binding; however, this region is crucial for the biological activity of auxins ^44^. Essentially, the molecule is no longer able to fit properly into the binding pocket, or the pocket does not allow the IAA to exist as a complete molecule within it, or both. The electronic binding energies between ABCG36 residues of the central pocket and the ligands IBA, CLX and IAA are -19.23, -8.11 and 34.08 kcal/mol, respectively. The positive energy for IAA essentially rules out IAA as an ABCG36 ligand that can be transported by ABCG36.

The 1D free energy surface (FES) of IBA translocation calculated from metadynamics simulations along the selected reaction coordinate (collective variable, Fig. 4b) exhibits an overall downhill process with some barriers from the intra-to extracellular direction for both Wt and L704F mutant versions of ABCG36 (Fig. 4f). In contrast, the FES from Wt ABCG36 CLX simulations exhibits low values at the intracellular ends of helices (around d_z_ of -1 nm), suggesting that this molecule can engage with this protein conformation easier than IBA (Fig. 4f-g). The FES for CLX from ABCG36^L704F^ simulations revealed a slightly elevated energy barrier between -1 and -0.5 nm in the L704F mutant compared to Wt, suggesting a more energy intensive access of CLX to the central pocket (Fig. 4g)

Since the FES around the extracellular gate (> 1.8 nm) was not conclusive about the role of this region in the transport, we also performed pulling simulations to measure the forces required to extract the compounds from the central binding pocket through the extracellular gate (Suppl. Fig. 5). The higher forces observed in the case of IBA removal from its initial position compared to CLX pulling in Wt ABCG36 (Fig 4i-j) agree with the DFT binding energy calculations. The elevated forces along the pathway observed in CLX pulling simulations for ABCG36^L704F^ indicated intensified interactions between CLX and the protein (Fig 4j), possibly contributing to diminished CLX transport in the mutant. Importantly, IAA, which is structurally very similar to IBA (Fig. 4a) but is not an ABCG36 substrate (Fig. 1 ^13,14,30^), also shows low energetic barrier at the beginning of the translocation pathway and very high barriers in the region of the central binding pocket of ABCG36 (Fig. 4h).

In conclusion, our simulations highlight that substrate translocation is likely controlled by different regions, including the entrance region (Fig 4f-g), the central binding pocket (Fig. 4h), and also the extracellular gate (Fig 4j). Interestingly, mutation in the extracellular gate can affect the energetics of the substrate interaction with distant protein regions (e.g. beginning of the translocation pathway, 1-2 nm from the extracellular gate, Fig. 4g), in an allosteric manner.

### L704 is a family-specific key residue of the extracellular gate

The di-leucine motif in ABCG2 is important for function but its sequence was found to be not conserved in human ABCGs (Suppl. Fig. 7a; ^22^). To get a deeper insight into the plant evolution of the key residues of the extracellular gate, we employed a recently published library of 1853 plant, full-size ABCG transporters extracted from the 1KP project ^25,45,46^. As for human ABCGs, the “leucine valve” is only poorly conserved in Arabidopsis and other plant ABCGs (Fig. 5; Suppl. Fig. 7): the first leucine following the conserved glycine is in most cases substituted by a phenylalanine (F703 in ABCG36), with an overall **G F** x x S/P R/K x x consensus motif (Fig. 5a). Overall, the extracellular gate of TH5 is slightly less conserved than that of TH11 (Fig. 5a-b).

We emphasized our analyses on the conservation of ABCG36 residues F703/L704 and F1374/F1375 as they are crucial residues of the extracellular gate limiting the translocation channel and are in close contact to the bound substrates, IBA and CLX (Fig. 5c; ^22,26,27^). We hypothesized that variations in the recognition of diverse substrates in various plant ABCG proteins might be reflected by their degree of conservation. We found that L704 and F1375 that are located most centrally in the translocation chamber (Suppl. Fig. 7b), revealed a far lower degree of conservation than F703 and F1374 (Fig. 5c; Suppl. Fig. 7b). While for F703 and F1374 of Arabidopsis ABCG36 in most cases (> 90%) the aromatic but hydrophobic phenylalanine is preserved, in other plant species L704 and F1375 are substituted by a limited choice of non-aromatic, hydrophobic amino acids, predominantly isoleucine, leucine and valine. Interestingly, in direct comparison, F1375 reveals a slightly higher level of variation than L704, the former also employing methionine and threonine (Figs. 5a-c).

Since higher plants have an advanced degree of chemodiversity provided by their specialized metabolism allowing for a more sophisticated lifestyle, including development and defense reactions ^8^, we compared the conservation of these key residues in the *Brassicaceae*, *Fabaceae,* and *Solanaceae* families containing beside Arabidopsis a series of important crop plants, like cabbage variations, soybean and tomato. While the overall degree of high and low conservation for F703/ F1374 and L704/ F1375, respectively, did not greatly differ amongst the families (Fig. 5d), we found a significantly higher and more equally distributed variability for L704 in the *Brassicaceae* family with a shift toward leucine and valine on the cost of phenylalanine. A similar shift toward leucine was also found for F1375.

Together with our transport and metadynamics data, these data support the evolution of L704 (and F1375) as *Brassicaceae* family-specific key residues of the extracellular gate that controls the identity of substrates during their transit from the substrate-binding cavity to the upper cavity.

### L704Y mutation in ABCG36 leads to a broadening of substrate specificity toward indolic compounds

In order to further challenge this apparent quality control function of the extracellular gate, we further investigated the transport specificity of L704 mutated versions of ABCG36 by site-directed mutagenesis. For choosing relevant mutations, we employed an amino acid property approach ^47^. Interestingly, like L704F all tested mutations of L704 retained IBA transport capacities, while this was not the case for CLX export (Fig. 6a-b). Exchange of leucine against alanine and positively and negatively charged arginine and aspartate, respectively, also abolished CLX export, while substitution for the aromatic phenylalanine or tyrosine did not (Fig. 6b).

**Figure 6.**
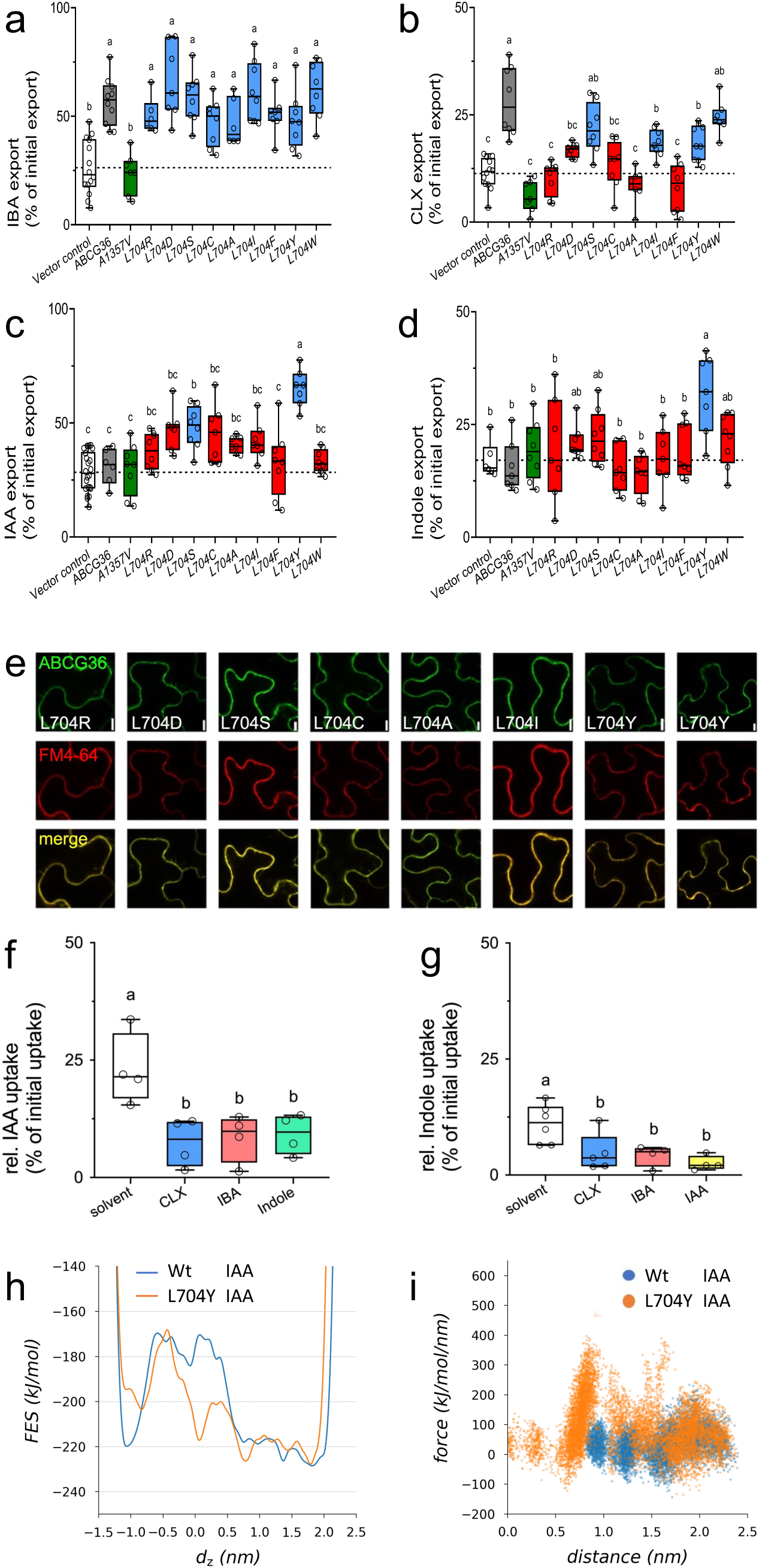
L704Y mutation in ABCG36 widens the transport selectivity to indolic compounds. **a-d.** IBA (**a**) CLX (**b**), IAA (**c**), and indole (**d**) export from *N. benthamiana* protoplasts after transfection with indicated mutant versions *ABCG36*. Significant differences (*p* < 0.05) of means ± SE (n ≥ 7 independent protoplast preparations) were determined using Brown-Forsythe and Welch ANOVA and are indicated by different lowercase letters. Additionally, means of mutant ABCG36 that are significantly different from Wt ABCG36 (grey fill) are indicated in red, while non-significant ones are in blue. ABCG36^A1357V^ (green) is included as a negative control. Significant differences (*p* < 0.05) of means ± SE (n ≥ 7 independent protoplast preparations) were determined using Brown-Forsythe and Welch ANOVA and are indicated by different lowercase letters. **e.** Confocal imaging of GFP-tagged versions of mutated *ABCG36* after transfection of *N. benthamiana* leaves. Short treatment of FM4-64 was used as PM markers; bar, 50 μm. Note that slightly enhanced PM signals of some mutant versions of ABCG36 in comparison to Wt are also found in respective FM4-64 controls and thus do not reflect higher expression levels; bar, 10 μm. **f-g.** Competition of IAA (**f**) and indole (**g**) uptake into microsomes prepared from ABCG36^L704Y^ transfected tobacco leaves; concentration of indicated competitors was 100x higher than radiolabeled IAA and indole. Significant differences (*p* < 0.05) of means ± SE (n ≥ 4 independent transport experiments) were determined using One-way ANOVA and are indicated by different lowercase letters. **h.** Free energy surfaces (FES) were calculated from metadynamics simulations with complexes of IAA with Wt and L704F ABCG36, respectively. **i.** Pulling simulations of IAA from the central binding pocket to the extracellular space were performed in simulations with Wt and L704Y ABCG36, respectively. Force distributions are plotted.

We included in our transport analyses also IAA and indole because they are, first, analogous to IBA or contain the indole core of all tested substrates and, second, they were neither transported by Wt nor L704F versions of ABCG36 (Fig. 6c-d; Suppl. Fig. 2e). Remarkably, we found a gain-of-IAA transport for polar but uncharged serine and the hydrophobic aromatic residue, tyrosine, but not for the closely related tryptophane (Fig. 6c). Interestingly, L704 exchange to tyrosine was the only mutation that enabled indole transport (Fig. 6d). Importantly, all mutations of L704 (except L704A) did not alter transport of the organic acid, BA (Suppl. Fig. 2e), commonly used as a diffusion control ^12^. Further, all mutations resulted in stable proteins on the PM and expression levels were comparable (Fig. 6e) allowing us to conclude that altered transport capacities were the direct cause of these mutations.

Next, we aimed to exclude that L704Y exchange did result in an uncoupling of ATPase activity and transport leading eventually to a permanently open conformation and thus an unspecific transporter. To do so, we performed competition experiments of IAA and indole transport using inside-out microsomes prepared from tobacco leaves infiltrated with ABCG36^L704Y^. As shown before for Wt ABCG36, transport of IBA and CLX ^14^, IAA and indole transport was completely abolished by 100-fold access of CLX, IBA, or indole/IAA, respectively (Fig.6f-g), indicating that ABCG36^L704Y^ is a functional ABC transporter.

The FES for IAA from metadynamics simulations with ABCG36^L704Y^ showed a reduced barrier near 0-0.5 nm compared to Wt (Fig. 6h). However, it remained high in the -1 to -0.5 nm region, suggesting that FES in the entry region may not entirely capture the molecular engagement with the translocation pathway, possibly occurring prior to the ATP-induced closure of intracellular transmembrane helices. Intriguingly, force distributions in ABCG36^L704Y^ pulling simulations for IAA, similar to those observed for non-substrate CLX in the L704F variant, were noted along the exit path (Fig. 6i). These findings imply that predominantly substrate access to and interactions within the central binding pocket govern ABCG36 transport. This conclusion is further supported by the rapid movement of IAA in an equilibrium simulation with Wt ABCG36, where this non-substrate molecule promptly moved out from the high-FES region of the central binding pocket and even translocated through the extracellular gate within 10 ns and did not exhibit high exit forces (Fig. 6i and Suppl. Fig. 6).

In summary, this mutational work further supports the conclusion that L704 is a key residue of the extracellular gate that provides quality control contributing to substrate specificity of ABCG36 toward a few indolic compounds. Exchange of the small, hydrophobic leucine to the hydrophobic but aromatic tyrosine broadens the substrate specificity slightly and allows for the export of structurally related, IAA and indole.

## Discussion

Despite the growing number of reported ABC transporter substrates and structures, the mechanisms underlying substrate identification and differentiation remain widely unclear. Intensive mutational analyses of TH residues that are thought to form substrate-binding sites and translocation pathways of paradigm ABC transporters, including human ABCB1, ABCC1 and ABCG2, and yeast PDR5, were shown to significantly influence the affinity and function of drug binding ^1,5,21,48–50^. Based on ABCG2 structures, residues L554 and L555 were suggested to form an extracellular “leucine plug”, for ABCG-type transporters, conceptualizing an additional layer of substrate specificity control: the opening and closing of the ‘leucine plug’ may act as a checkpoint during the transport reaction. Khunweeraphong et al. (2019) extended this concept of a symmetric pump gate to six hydrophobic residues forming a “valve-like structure”, contributing to the regulation of substrate efflux potentially trading off translocation speed with substrate selectivity (Manolaridis et al., 2018).

In this work we further hardened but also widely extended this overall concept by taking advantage of the recently published Arabidopsis *ABCG36* allele, *abcg36-5*, that contains a L704F point mutation in the extracellular gate and for which an uncoupling of ABCG36 functions in IBA-stimulated root growth from ABCG36 activity in extracellular defense was suggested ^29^. Using heterologous and homologous transport systems (Figs. 1-2) we clearly demonstrate that this uncoupling of ABCG36 function in growth from defense is due to a unilateral loss-of CLX transport capacity, while IBA transport was preserved. This contrasts with the *abcg36-6* allele that contains an A1357V substitution in the residue, which is a part of TH11, but not part of the extracellular gate. Importantly, in contrast to the T-DNA insertion mutant (null) allele, *abcg36-4,* used here as a negative control, both ABCG36^L704F^ and ABCG36^A1357V^ are expressed at similar levels on the PM (Figs. 1g and 2e, ^29^).

ATPase activities (Fig. 1h) of ABCG36^L704F^ were comparable to Wt ABCG36 and showed likewise a stimulation by IBA and CLX, while ABCG36^A1357V^ were slightly enhanced but lacked substrate stimulation. This indicates that unlike ABCG36^L704F^ mutation in ABCG36^A1357V^ resulted most likely in an uncoupled, transport-incompetent transporter. Interestingly, in contrast to L555A exchange in ABCG2 resulting in lowered substrate-stimulated ATPase activity but twice the translocation activity of the wild-type (Manolaridis et al., 2018), the L704F mutation in ABCG36 preserved ATPase activity and unilateral transport capacity.

Metadynamics simulations of Wt and L704F mutant versions of ABCG36 (Fig. 4) using an AlphaFold2-predicetd structure in a closed conformation ^43^ provided a FES- and force-based interpretation for the transport data, indicating why IBA is a substrate for both Wt and ABCG36^L704F^, while CLX is only for Wt ABCG36 (Figs. 1-2). Differences in the FES profiles around the central binding pocket and pulling forces towards the valve region (Fig. 4) suggest that the export of different substrates is determined by various regions of the transport pathway.

Metadynamics simulation with the non-substrate IAA ^13,14,30^ revealed high barriers for IAA movement around the central pocket (Fig. 4h). These barriers explain the remarkable ability of ABCG36 to discriminate between the structurally very similar IBA and IAA (Fig. 4a). DFT analyses provided evidence that this high barrier IAA might be caused by drastic deformation of the electronic structure after binding to the ABCG36 binding pocket (Fig. 4d-e). This change in the electron system results in a positive binding energy preventing transport. This capability of ABCG36 is apparently essential because the IAA precursor-IBA is thought to function in overlapping but distinct developmental pathways ^51^.Our simulations with IBA and CLX indicate that multiple mechanisms could underlie substrate recognition, such as the entry to the central binding pocket, interactions within this pocket, and interactions around the extracellular gate. However, the increased necessary forces for both the non-substrate CLX and the IAA substrate for ABCG36^L704F^ and ABCG36^L704Y^, respectively, weaken the role of direct molecular interactions around the extracellular gate in substrate discrimination. These observations strengthen the role of the central binding pocket in substrate recognition, underlined by the presence of non-transported molecules within this pocket in cryo-EM structures ^48^. In summary, while mutations in the extracellular gate could alter interactions with various molecules, our *in silico* findings downplay these direct effects on transport. Instead, they highlight the importance of allosteric interactions between the valve residues and the central binding pocket, influencing substrate recognition and transport.

An evolutionary analysis uncovered that like for human ABCGs, the extracellular gate is poorly conserved in Arabidopsis and other plant ABCGs (Fig. 5; Suppl. Fig. 6). L704 and F1375, which are in closest vicinity to the transported substrates and that are located most centrally in the translocation chamber (Fig. 4c; Suppl. Fig. 7b) revealed a far lower degree of conservation than F703 and F1374 (Fig. 4c). A comparison of residue conservation in the major crop families uncovered, in contrast to the overall degree of high and low conservation for F703/ F1374 and L704/ F1375, respectively, which does not greatly differ amongst the families (Fig. 5d), a more equal distribution and variability for L704 in the *Brassicaceae* family but not for F1375. A reasonable explanation is that an enhanced variability at this position has allowed ABCG36-like transporters in the *Brassicaceae* family to accommodate several indolic substrates, such as CLX and IBA, crucial for defense and development, respectively.

Mutational work further supported the conclusion that L704 is a key residue of the extracellular gate that provides quality control contributing to substrate specificity of ABCG36 toward a few indolic compounds. All tested mutations of L704 (including L704F and L704A) preserved IBA transport capacities, while this was not the case for CLX export (Fig. 6a). This might suggest that most likely the additional aromatic ring of CLX allows for a discrimination by using a contact with L704. Remarkably, exchange of L704 against the aromatic residue, tyrosine, but not for structurally similar tryptophane or phenylalanine resulted in a gain-of-IAA and indole transport (Fig. 6). Competition experiments excluded that L704Y mutation did result in an uncoupling of ATPase activity and transport leading to a permanently open conformation and thus an unspecific transporter (Fig. 6f-g).

This broadening of substrate specificity by L704Y but not L704F or L704W mutations is remarkable because all three aromatic residues are chemically and structurally similar; e.g. phenylalanine and tyrosine amino acids differ just by a hydroxyl group.

Therefore, we conclude that the change of a leucine to the aromatic phenylalanine limited transfer through the extracellular gate, while further addition of a hydroxyl group widened substrate specificity. Interestingly, tyrosine is absent in any taxa of higher plants for the L704 position, but it is frequent for positions of F703 and 1374 (Fig. 5) and is part of the so-called “apoplastic gate” in the ABA exporter ABCG25 (Y565; ^27^).

Taken together, our transport, metadynamics simulation and evolutionary work support the evolution of L704 as a *Brassicaceae* family-specific key residue of the extracellular gate that may control the translocation process in different regions, allosterically at distant parts based on simulations and directly indicated by experiments. Initially, we aimed to correlate the composition of the extracellular gate with known substrate specificities of plant PDRs (including IBA, CLX, and ABA), but this trail failed. Also, data from metadynamics simulations did not reveal large differences in FES around the extracellular gate of various compound/protein complexes, except in the case of IBA. As such we believe that the gate does not exclusively contribute to substrate specificity but that it instead may provide a final quality check of chemically closely related molecules, especially for the case that a non-substrate was falsely recognized in the central binding pocket region.

From the plant’s perspective, such a final quality control on the substrate level provided by the extracellular gate of ABCG36 is of major importance because as mentioned above, both ABCG36 substrates, IBA and CLX, serve as crucial substrates in two separate programs, growth and defense, respectively. While the need for such a final quality control is obvious for targeted CLX export provided by ABCG36 expressed toward the attacking pathogen ^36^, the opposite scenario might be also critical. A physiological consequence of prolonged defense is growth inhibition ^16,52^ because essential resources are thought to be allocated toward defense ^16^. On the molecular level, this coordination of substrate preferences on ABCG36 was shown to be provided by phosphorylation situated in the linker (mainly S825 and S844) repressing IBA export and allowing for CLX export and thus defense ^14^. Currently, it is unknown if these phosphorylation events exert an effect on the central binding pocket or extracellular gate, however, this option is not unlikely, as linker phosphorylation-sites and L704 of the extracellular gate are directly connected by the THs of TMD2 (not shown).

In summary, our findings support the concept of a gate-keeper function of the extracellular gate and therefore contribute to our understanding of how ABCG substrate specificity is encoded. This is of interest because many ABCG proteins are also of clinical relevance, since their promiscuous substrate selectivity causes pleiotropic or multidrug resistance phenomena ^1^. This is most obvious for the direct involvement of ABCGs in several genetic diseases (like ABCG5/G8 in sterol sitosterolemia or ABCG2 in gout) or chemotherapy failure (ABCG2) but also the treatment of fungal pathogens, like *Candida* species ^1,53^. Infections by Candida, themselves containing a plethora of ABCG transporters, are increasingly providing a threat to immunosuppressed people ^53^.

## Acknowledgments

We would like to thank L. Charrier (UNIFR) and U. Smolka (IPB) for excellent technical assistance, J. Huyhn (UNIFR), and T. Romeis (IPB) for experimental support. This work was supported by grants from the Swiss National Funds (project 31003A_165877 and 310030_197563 to MG and project 310030_184769 to CSR), the *Pool de Recherche* of the University of Fribourg (to MG), the Polish National Science Centre (grant 2017/27/B/NZ1/01090 to MJ and 2021/41/N/NZ1/04030 to MJ and KP), the German Research Foundation (grant RO 1172/6−1 to SR), National Research, Development and Innovation Office NKFIH-137610, TKP2021-EGA-23 (to TH) and the China Scholarship Council (to JX).

## Author contribution

MG, TH and JX designed research; JX, AS, OT, JB, KP, JZ, NF, and TH performed research; JX, AS, OT, JB, KP, JZ, SR, NF, TH, and MG analyzed data; MJ, SR, TH and MG supervised work; MG wrote the manuscript, all authors commented on the manuscript.

## Methods

### Plant material

The *following Arabidopsis thaliana* lines in ecotype Columbia (Col Wt) were used: *abcg36-4/pen3-4* (SALK_000578, Stein *et al.* 2006), *abcg36-6/pen3-6/pdr8-115* (Strader *et al.* 2009). gl1 (gl1 Wt) was used as the wild-type control for *abcg36-5/pen3-5* ^29^.

Mutant versions of *ABCG36* were obtained by QuickChange (Agilent Company, USA) site-directed mutagenesis of *35S:ABCG36* (Kim *et al.* 2007), *35S:ABCG36-GFP* ^14^and *ABCG36:ABCG36-GFP* (Stein et al., 2006). Mutated plasmids were used to transform *abcg36-4* by floral dipping to generate *ABCG36:ABCG36^L704F^*-GFP (*abcg36-4*) #1, *ABCG36:ABCG36^L704F^*-GFP (*abcg36-4*) #2, *ABCG36:ABCG36^A1357V^*-GFP (*abcg36-4*) #1, *ABCG36:ABCG36^A1357V^*-GFP (*abcg36-4*) #2, *ABCG36:ABCG36^L704F^ ^A1357V^*-GFP (*abcg36-4*) #1 and *ABCG36:ABCG36^L704F^ ^A1357V^*-GFP (*abcg36-4*) #2. Isogenic, homozygous lines for the transgene in the F3 generations were used for further analyses.

### Plant growth conditions

Seedlings were generally grown on vertical plates containing 0.5 Murashige and Skoog media, and 0.75% phytoagar at 12h light per day. Developmental parameters, such as primary root lengths, were quantified as described in (Ruzicka et al., 2010). All experiments were performed at least in triplicate with 30 to 40 seedlings per experiment.

### *Fusarium oxysporum* elicitor treatments and root infection assays

*Fusarium oxysporum f. sp. conglutinans* (Fo), strain 699 (Fo699; ATCC 58110), and strain 5176 (Fo5176) were grown as described in ^54^. The fungal elicitor mix was prepared from Fo5176 as described elsewhere ^55^. *F. oxysporum* infection assays of Arabidopsis roots in soil were performed with strain Fo699 as reported in ^54^. In short, 3-week-old *Arabidopsis thaliana* plants were infected by pipetting 10 ml conidia suspension (10^7^ conidia/ml) directly into the soil contained in a 125 ml plastic pot harboring a single plant. Fresh weight and disease sensitivity scores were obtained by measuring rosette weight and by observing chlorotic and necrotic symptoms scored at a scale from 1-10 as number of affected leaves per plant, respectively, two weeks after inoculation. 15-30 plants were employed per genotype per experiment in each of three independent experiments (n = 3).

### Phytophthora infestans infection assays

Rosette leaves of 5-week-old *A. thaliana* plants were drop-inoculated with zoospore suspensions of *P. infestans* Cra208m2 ^56^ or of *B. cinerea* B05.10 by applying 10 µl drops containing 5 x 10^5^ or 5 x 10^4^ spores ml^-1^, respectively, onto the adaxial leaf surface. Thirty drops each were collected and immediately frozen in liquid nitrogen for storage. For extraction, the drops were evaporated to dryness, and samples were dissolved in 60 µl of 30% methanol, sonicated for 15 min, and analyzed by LC-MS. Inoculation of seedlings was performed as described (He et al. 2019). Arabidopsis seedlings (Col-0, *abcg36-6*, Gl1 Wt and *abcg36-5*) were grown in 2 ml liquid medium (1/2 strength MS with sucrose) for 12 days. One day before inoculation with *B. cinerea* (5 x 10^3^ ml^-1^) or *P. infestans* zoospores (1 x 10^5^ ml^-1^), the medium was removed and replaced by 2 ml 1/2 MS without sucrose. 600 µl of the medium were collected 24h post-inoculation, evaporated to dryness, and re-dissolved in 120 µl of 30% methanol. CLX levels were determined by UPLS-ESI-QTOF-MS.

### Transport studies

^3^H-Indole (ARC ART2269, 25 Ci/mmol), ^3^H-IAA (ARC ART0340, 25 Ci/mmol), ^3^H-IBA (ARC ART2533, 10 Ci/mmol), ^14^C-BA (ARC ART0186A, 55 mCi/mmol) and ^3^H-CLX (3-thiazol-2-yl-indole; custom-synthesized by ARC, 56 Ci/mmol) export from *Arabidopsis* and *Nicotiana benthamiana* mesophyll protoplasts was analyzed as described ^39^. *N. benthamiana* mesophyll protoplasts were prepared 4 days after agrobacterium-mediated transfection of combinations of indicated plasmids or plasmid combinations. Transfections with an anti-silencing strain, RK19, were used as vector control. Relative export from protoplasts is calculated from exported radioactivity taken at t = 0, 5, 10, 15 and 20 min. as follows: (radioactivity in the supernatant at time t = x min.) - (radioactivity in the supernatant at time t = 0)) * (100%)/ (radioactivity in the supernatant at t = 0 min.); presented are mean values from > 4 independent protoplast preparations at t = 15 min.

^3^H-IAA and ^3^H-Indole uptake into *Arabidopsis* vesicles prepared from Arabidopsis lines grown as mixotrophic liquid cultures was measured in the absence (solvent) or presence of 1000 x access of nonlabelled Indole, IAA, IBA, or CLX, respectively, as described in ^30^.

### Measurement of ATPase activity

ATPase activity was measured from microsomes (0.06 mg/ml) prepared from tobacco plants transfected with vector control or Wt and mutant versions of *35S:ABCG36-GFP* ^39^ using the colorimetric determination of ortho-phosphate released from ATP ^57^. Briefly, microsomes were added to ATPase buffer (20 mm MOPs pH 7.7 or pH 9.0, 8 mm MgSO_4_, 50 mm KNO_3_, 5 mm NaN_3_, 0.25 mm Na_2_MoO_4_, 2.5 mm phosphoenolpyruvate, 0.1% pyruvate kinase) in the presence and absence of 0.5 mM sodium ortho-vanadate. The reaction was started by the addition of 15 mm ATP and incubated at 37°C for 15 min with shaking. The amount of Pi released in the absence (solvent control) or presence of IBA, IAA, CLX, indole, or *ortho*-vanadate (50 μM) was quantified using a Cytation 5 reader (BioTek Instruments).

### Metabolite profiling using LC*-MS*

Untargeted metabolite profiling was performed as described ^30^. Targeted relative quantification of metabolites was done on the peak areas of characteristic extracted ion chromatograms in Bruker’s QuantAnalysis V4.1 software (Bruker Daltonik).

### Confocal laser scanning

For imaging, seedlings were generally grown for 5 days on vertical plates containing 0.5 Murashige and Skoog media, 1% sucrose, and 0.75% phytoagar at 16h (long day) light per day. For chemical treatments, seedlings were transferred for 12h on test plates containing the indicated chemicals or solvents. For confocal laser scanning microscopy work, an SP5 (Figs. 1g, 2e, 6e, 7e) confocal laser microscope was used. Confocal settings were set to record the emission of GFP (excitation 488 nm, emission 500-550 nm), and FM4-64 (excitation 543 nm, emission 580-640 nm). Images were quantified with ImageJ (http://imagej.nih.gov/ij/) using the Fiji plugin and identical settings for all samples. As regions of interest either whole images or individual cells marked using the freeform tool were used for quantification; these are indicated in the legends.

### Evolutionary PDR analyses

To select full-size ABCG from multiple groups of plants, we subjected One Thousand Plant Transcriptomes (1KP) data (https://db.cngb.org/onekp/) ^45,46^ to tBLASTn searches using ABCG36 as a query sequence with an E-value cutoff of 1e−5. Obtained sequences were further filtered as described in ^25^. One thousand eight hundred and fifty-three 1KP samples were assigned to a full-size ABCG subfamily and used for the WebLogo analysis ^58^. Multiple sequence alignment (MSA) of the full-size ABCG amino acid sequences was performed using the MUSCLE algorithm^59^. To analyze the amino acids occurrence rate in selected residues from the full-size ABCG sequences, firstly BLASTp search was performed independently for *Brassicaceae, Fabaceae,* and *Solanaceae*, using the sequence of AtABCG36 as a query with an E-valu cut-off of 0.05. Obtained sequences were then filtered by length (between 950 and 1800) and checked to contain plant PDR signature motives ^60^ (LLXGPP, GLXSS, GLDARXAAXVMR, VCTIHQP). 598 sequences from *Brassicaceae*, 762 from *Fabaceae*, 450 from *Solanaceae,* and 1599 sequences from all taxa (obtained from 1KP) were aligned using the MUSCLE algorithm and analyzed using Python.Protein structures and chemical formulas

The AlphaFold2 predicted ABCG36 structure (AF-Q9XIE2-F1) was used (PMID: 34265844; PMID: 34791371). ATP molecules and Mg^2+^ ions were inserted after the structural alignment of nucleotide-binding domains of this ABCG36 structure and our formal ABCG2 homology model (PMID: 32979053). Regions with high uncertainty (residues 1-39 and 809-863) were deleted since they may exert unwanted effects in molecular dynamics simulations. IBA, IAA, IND, and CLX 3D structures were downloaded from PubChem (IDs: 8617, 802, 798, and 636970).

### *In silico* docking

The small molecules were docked to the extracellular (EC) side of the extracellular gate of the AF-predicted ABCG36 structure using AutoDock Vina (PMID: 34278794). The following options were set: grid size = 22.50 Å, center_x = 1.4, center_y = 0.22, center_z = 15, exhaustiveness = 64. The predicted pose with the lowest score/binding energy was used in MD simulations.

### Molecular dynamics simulations

The input files for all steps (energy minimization, equilibration, and production run) were generated by the CHARMM-GUI web interface (PMID: 18351591) by submitting the structure of a complex, including the protein, ATP, and a small molecule. CHARMM36m all-atom forcefield was used for the protein since it removes the bias towards left-handed helices and improves the dynamics of backbone dihedrals (PMID: 27819658). CHARMM, General ForceField (CGenff) parameterized IBA, CLX, and IAA (PMID: 19444816). Periodic box conditions were applied with a rectangular box, which had proportions sufficient to fulfill the minimum image convention (approx. 10-10 AL from every edge). The full-length structures were oriented according to the OPM database. Mutations were introduced via CHARMM-GUI. A mixed membrane bilayer was built, including 1:1 POPC: PLPC (1-palmitoyl-2-oleoyl-sn-glycerol-3-phosphocholine:1-pal-methyl-2-linoleoyl-sn-glycerol-3phosphocholine) in the extracellular leaflet and 37:43:12:8 POPC: PLPC: POPS: DMPI25 (POPS: 1-palmitoyl-2-oleoyl-sn-gglycerol3-phospho-L-serine, DMPI25: dimyristoyl-phosphatidylinositol 2,5) in the intracellular leaflet. The following additional options were adjusted: i) terminal residues were patched by ACE (acetylated N-terminus) and CT3 (N-methylamide C-terminus), ii) 150 mM KCl in TIP3 water was used, iii) grid information for PME (Particle-Mesh Ewald) electrostatics was generated automatically and iv) a temperature of 303.5 K was set. The structures were energy minimized using the steepest descent integrator (maximum number to integrate: 50,000 or converged when force is <1,000 kJ/mol/nm). From the energy-minimized structures, an equilibrium simulation was forked for 100 ns. During these simulations, a Berendsen thermostat and a Parinello-Rahman were applied for equilibration and production runs, respectively. Besides, Parinello-Rahman barostat with a semi-isotropic, 1 atm pressure coupling, a time constant of 5 ps in the plane of the membrane, with the compressibility of 4.5 x 10-5 bar^-1^, was applied (PMID: 19444816, PMID: 26631602). The cut-off value considered for the long-distance interactions was 12 AL. Electrostatic interactions were calculated using the fast, smooth PME algorithm, and the LINCS algorithm was used to constrain bonds at 1.2 nm. Constant particle number, pressure, and temperature ensembles were used with a time step of 2 fs. Simulations were performed using GROMACS 2022 (https://doi.org/10.1016/j.softx.2015.06.001).

Well-tempered metadynamics simulations were executed with GROMACS 2022 patched with PLUMED 2.8.1 (PMID: 31363226, https://doi.org/10.1016/j.cpc.2013.09.018). Multiple walker setups (n=4 or 8, depending on hardware) were used. The total simulation time was 2 µs for each complex (total time: 7 systems x 2 μs = 14 μs). CV was defined as the Z component of the distance between the center of geometry of ABCG36 residues in the central transmembrane helices (TH) (a.a 582-597: FGMIINMFNGFAEMAM, 685-700: IANTGGALTLLLVFLL, 1250:1265: LYAAIIFVGINNCSTV, 1356-1371:

VASIFASAFYGIFNLF). The parameters used were ARG=CV PACE=500 (1ps), HEIGHT=0.6, SIGMA=0.05, BIASFACTOR=10.0, TEMP=303.15K, GRID_MIN=-1.5 GRID_MAX=2.4, GRID_WSTRIDE=1000. The movements were constrained with LOWER_WALL (d_z_ = -1.1 nm) and UPPER_WALL (d_z_ = 2.0 nm) to inhibit the escape of the small molecules from the four helices that would have rendered the simulations unfeasible due to the extremely low probability of returning to the pathway. FES was calculated using the sum_hills tool of PLUMED.

For simulations pulling each compound from the intracellular to the extracellular side of the extracellular gate, the system was prepared as follows. First, PLUMED was used to pull the small molecule into the binding pocket, using the same CV as for Metadynamics simulations. The parameters used were ARG=CV, STEP0=0, AT0=INITIAL_DISTANCE, KAPPA0=1000, STEP1=4,500,000, AT1=0.2, STEP2=5,000,000, AT2=0.2, KAPPA2=0. Then, an unconstrained, short (10 ns) equilibrium simulation was performed, and the resulting trajectory was sampled for 25 frames with different velocities as input for pulling simulations. The pull-out was performed using the GROMACS pull code. Umbrella pulling was selected with the same CV as metadynamics; thus, the force was proportional to the displacement. Small molecules were pulled along the positive Z axis (Suppl. Fig. 5) with a force constant and velocity of 500 kJ/mol/nm and 0.0005 nm/ns respectively.

Analysis was done using MDAnalysis (PMID: 21500218) and Matplotlib (doi: 10.1109/MCSE.2007.55) Python packages. PyMOL Molecular Graphics System (Version 2.0 Schrödinger, LLC) was used for molecular visualization tasks.

### Quantum Chemical Modeling

The initial ABCG36 geometry (PMID: 34265844; PMID: 34791371) was derived from protein structure predictions using AlphaFold2 (AF-Q9XIE2-F1). The size of the binding pocket region was determined through docking methods using AutoDock Vina. Molecular dynamics were used to perform several geometric conjugations of a pocket of 630 atoms with each of the different ligands, using MM+ force fields and their respective pre-optimizations. At least two conformations pre-optimized at the MM+ level were taken for each ligand.

The following step involved obtaining optimizations using Density Function Theory methods, which included a Van der Waals D3 correction (DFT-D) with the bp86 functional and the double zeta polarized atomic basis set level [def -SV(P)]. After the calculation of each molecular component, the pocket-ligand binding energies were determined by first calculating the electronic interaction energy using the equation E_el_ = E_PL_ - (E_P_ + E_L_ ), where E_el_ is the electronic binding energy, E_PL_ is the electronic interaction energy between the pocket and ligand, E_P_ is the electronic interaction energy within the pocket, and E_L_ is the electronic interaction energy of the ligand. All calculations were performed with Turbomole (http://www.turbomole.com). The analysis of the deformation of the electronic structure and changes in the potential surface was carried out using the theory of deformation of atoms in molecules ^61^

### Statistical analyses

Data were statistically analyzed using Prism 10.1.1. (GraphPad Software, San Diego, CA).

## Data availability

This article does not contain any original code. Requests for data should be made to and will be fulfilled by M.M Geisler, provided the data will be used within the scope of the originally provided informed consent. Source data are provided with this paper.

**Supplemental information included in this work.**

**Supplementary Figure 1: In the *abcg36-5* allele, ABCG36-mediated IBA – but not camalexin - detoxification activity is retained**

**Supplementary Figure 2: Transport controls**

**Supplementary Figure 3: Structural comparison of ABCG2 and ABCG36**

**Supplementary Figure 4: Monitoring the convergence of metadynamics simulations**

**Supplementary Figure 5: Scheme of ABCG36 pulling simulations**

**Supplementary Figure 6: Non-substrate IAA exits in a short equilibrium simulation with Wt ABCG36**

**Supplementary Figure 7: Conservation of essential residues in the extracellular gate of plant PDR-type ABCGs**

**Supplementary Table 1: Primer sequences for QuikChange site-directed mutagenesis**

## Footnotes

1 In that respect, we would like to suggest to the community to use in the future only the term “extracellular gate” instead of similar or equivalent terms, like “apoplastic gate” (Huang *et al.,* 2023), “di-leucine valves” (Khunweeraphong *et al.,* 2019), “leucine gate” or “leucine plug” (Taylor et *al.,* 2017; Yu *et al.,* 2021). First, it appears that the conservation of leucines is very poor in these gates (see Suppl. Fig. 6), and second, the term “apoplastic” (referring to the extracellular space of plant cells, an equivalent of the extracellular matrix (ECM)) does not exist in non-plant organisms.

