## Supplementary material for "A key residue of the extracellular gate provides quality control contributing to ABCG substrate specificity": Suppl. figures 1-7

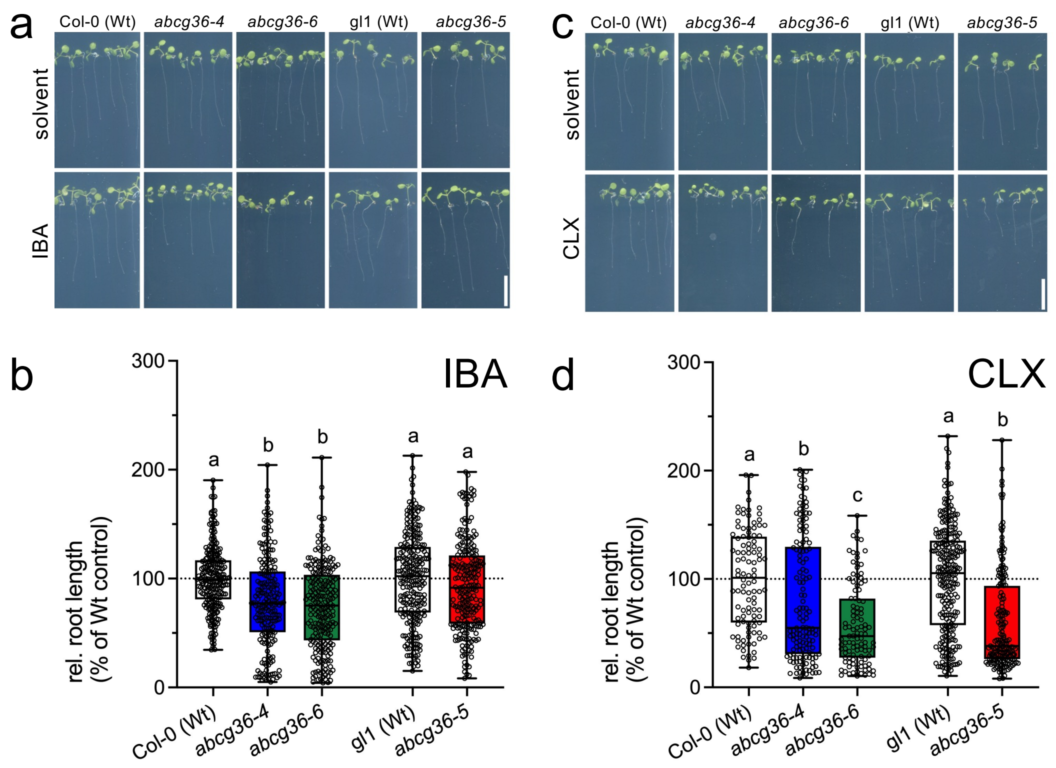


**Supplementary Figure 1: In the *abcg36-5* allele, ABCG36-mediated IBA – but not camalexin - detoxification activity is retained**

**(a-d)** Relative root length of indicated Arabidopsis lines grown in the presence of 7.5 µM IBA **(a-b)** and 5 µg/mL CLX **(c-d)** for 12 days. Wt (Col-0 and gl1) growth is set to 100%. Results are mean values (±SE) of 4 independent experiments with 10-20 seedlings each. Different lowercase letters indicate that the means are significantly different (Ordinary one-way ANOVA with a post hoc Sidak's multiple comparison test, $p < 0.05$). Scale bar = 1 cm.

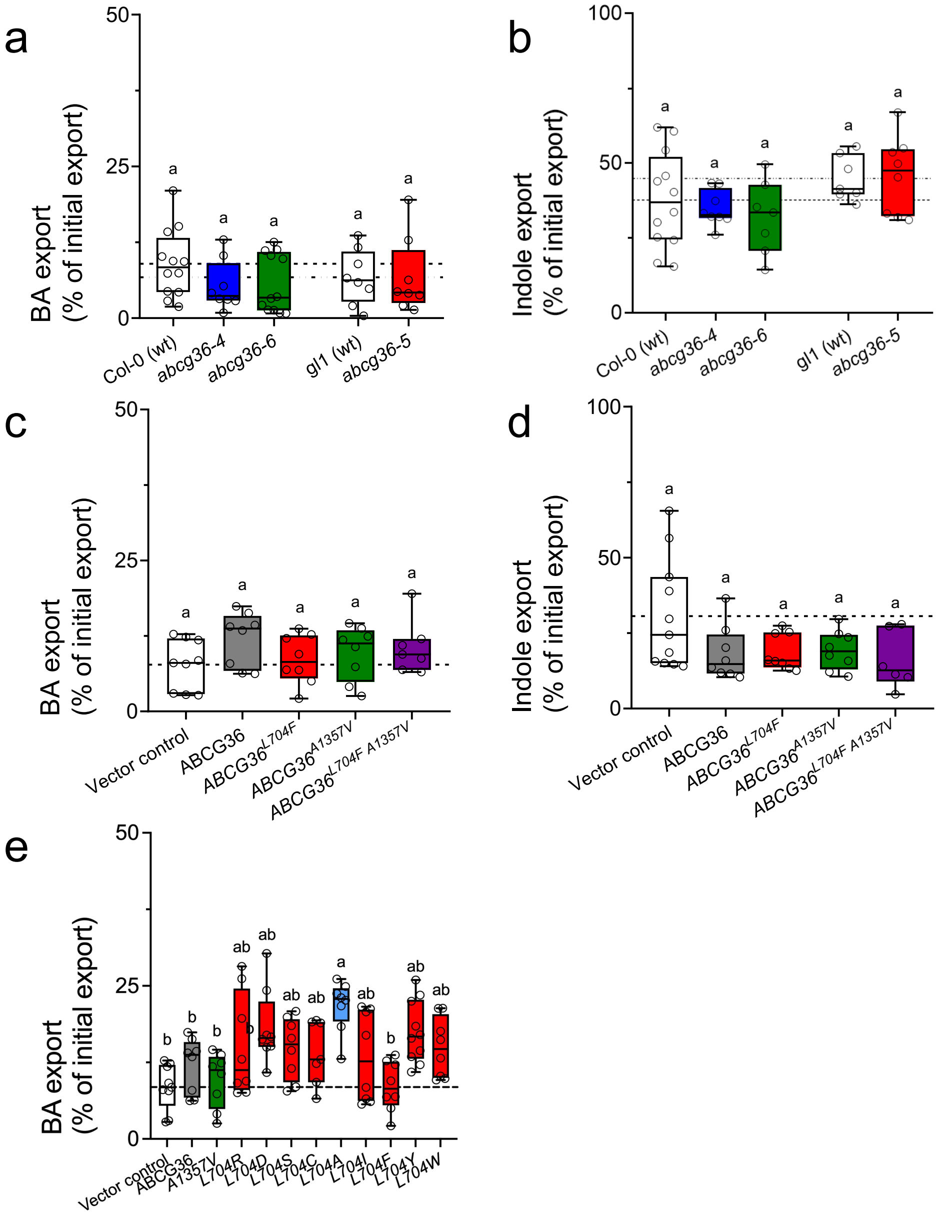


**Supplementary Figure 2: Transport controls**

**a-b.** BA (**a**) and indole (**b**) export from *Arabidopsis* protoplasts prepared from indicated *ABCG36* loss-of-function alleles. Significant differences ($p$ < 0.05) of means ± SE (n ≥ 7 independent protoplast preparations) were determined using Brown-Forsythe and Welch ANOVA and are indicated by different lowercase letters.

**c-d.** BA (**c**) and indole (d) export from *N. benthamiana* protoplasts after transfection with indicated mutant versions *ABCG36*. Significant differences ($p$ < 0.05) of means ± SE (n ≥ 7 independent protoplast preparations) were determined using Brown-Forsythe and Welch ANOVA and are indicated by different lowercase letters.

**e.** BA export from *N. benthamiana* protoplasts after transfection with indicated mutant versions *ABCG36*. Significant differences ($p$ < 0.05) of means ± SE (n ≥ 7 independent protoplast preparations) were determined using Brown-Forsythe and Welch ANOVA and are indicated by different lowercase letters. Additionally, means of mutant ABCG36 that are significantly different to Wt ABCG36 (grey fill) are indicated in red, while non-significant ones are in blue. ABCG36A1357V (green) is included as a negative control.

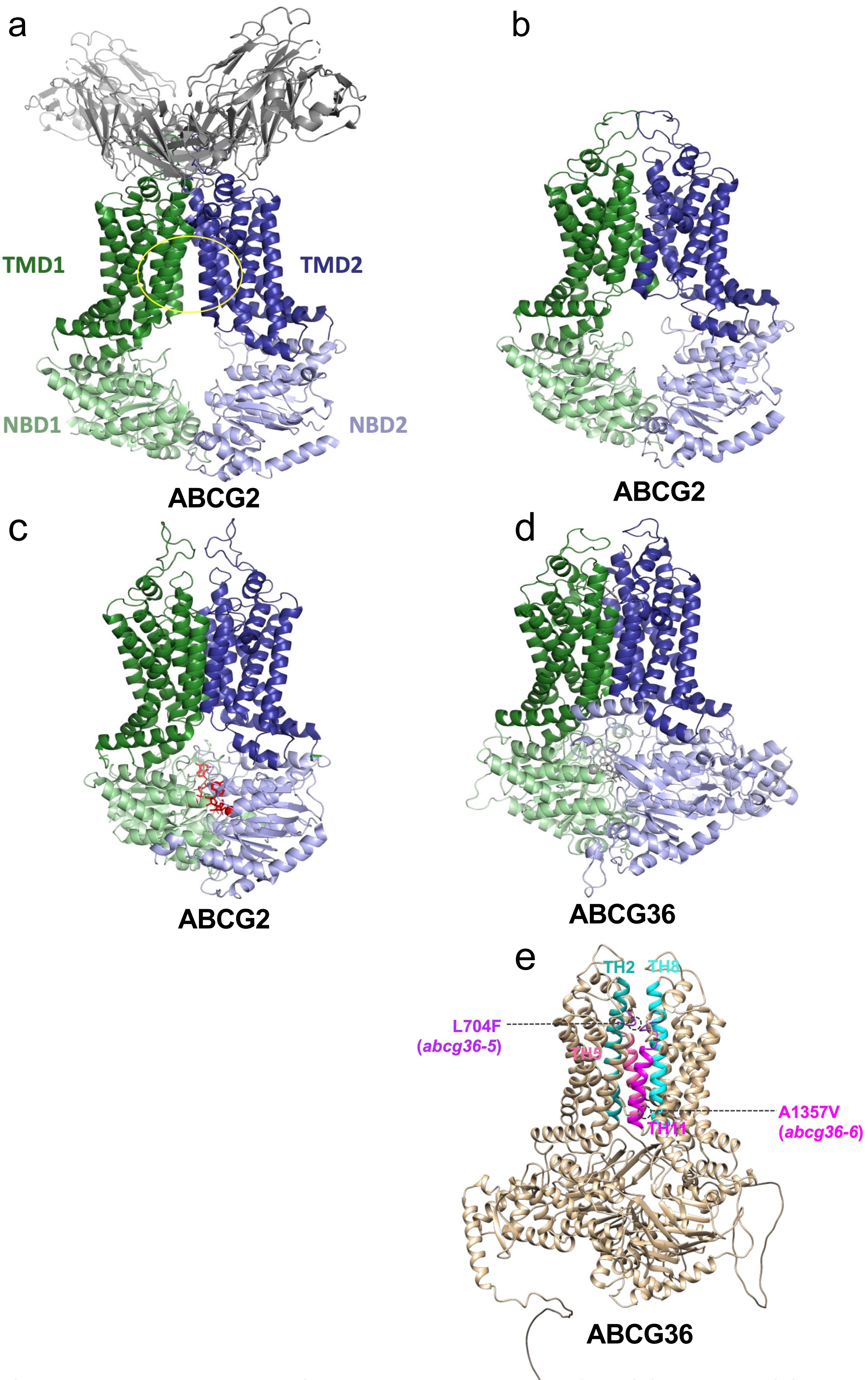


**Supplementary Figure 3: Structural comparison of ABCG2 and ABCG36**

**a.** The functional unit of ABC proteins consists of two transmembrane domains (TMD, dark green, and dark blue) and two nucleotide-binding domains (NBD, green and blue). The TMDs and NBDs of the human ABCG2 multidrug transporter are in an open conformation in the absence of ATP and with extracellular stabilizing antibody (gray); the central binding pocket is indicated by a yellow circle (PDBID: 5nj3).

**b.** In an *apo* ABCG2 structure (PDBID: 6vxf), the NBDs are open but TMDs are closed.

**c.** In the presence of ATP both NBDs and TMDs are closed (PDBID: 6hbu).

**d-e.** The AlphaFold2-predicted ABCG36 structure exhibits a closed conformation. This structure was used in our study, since this structure is the closest to the transport-competent conformation (**d**). Location of employed ABCG36 alleles, *abcg36-5* (L704F) and *abcg36-6* (A1357V), respectively, as well as THs are illustrated (**e**).

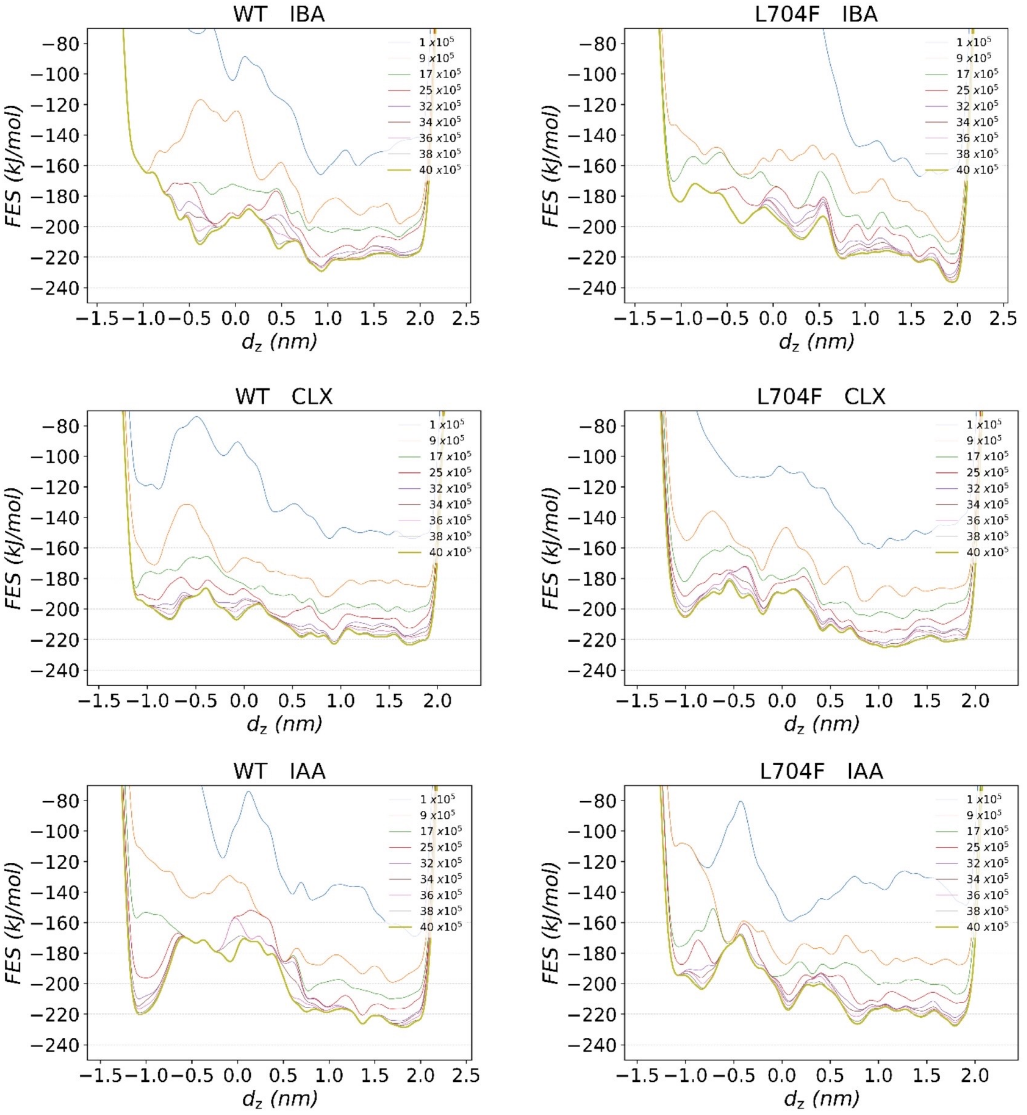


**Supplementary Figure 4: Monitoring the convergence of metadynamics simulations.**
FES for Wt ABCG36 (Wt) and indicated L704 of ABCG36 were calculated and plotted at indicated strides (every $n^{th}$ Gaussian kernel) using the sum_hills command of PLUMED.

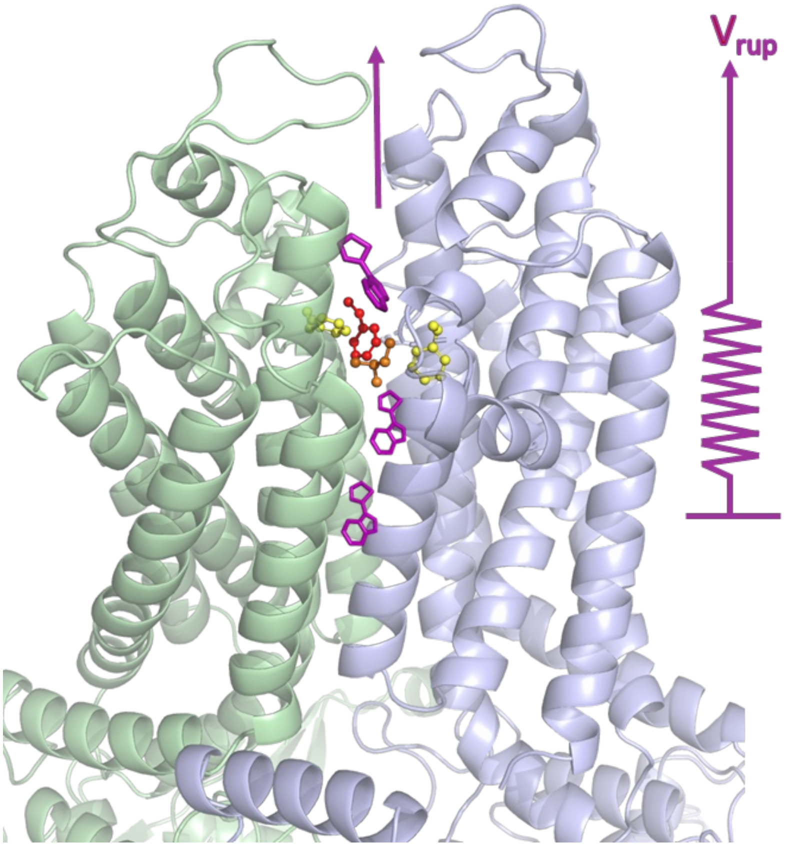


**Supplementary Figure 5: Scheme of ABCG36 pulling simulations**
Force-constrained GROMACS umbrella pulling was used to displace the molecule from the central binding pocket to the extracellular space. A harmonic potential was applied ($v_{rup}$: the velocity at which the harmonic potential was retracted). Orange, L704; yellow, F703 and F1374; red, F1375.

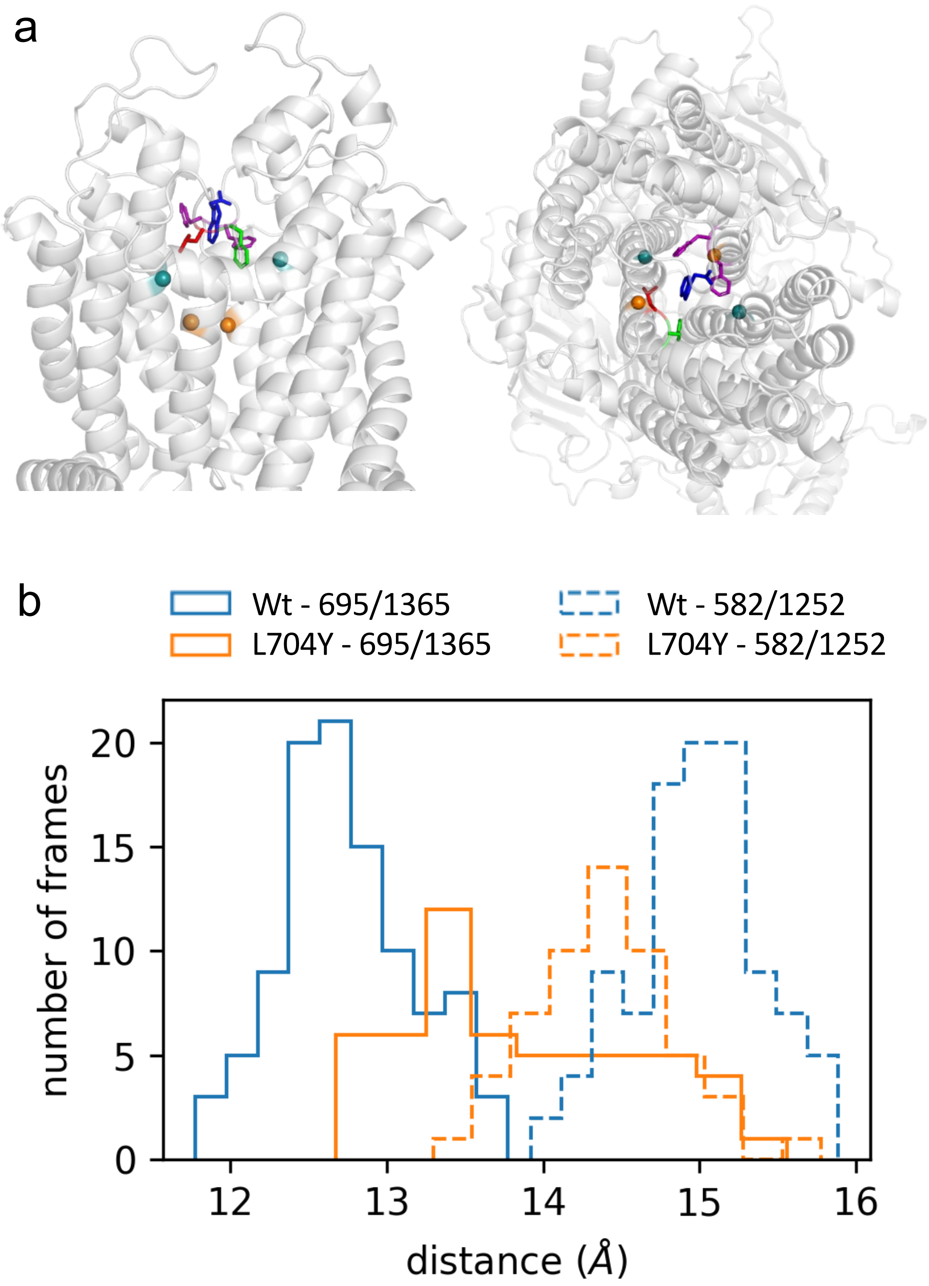


**Supplementary Figure 6: Non-substrate IAA exits in a short equilibrium simulation with Wt ABCG36**

**a.** IAA passes through the extracellular gate in 10 ns without applying any bias. ABCG36 TMDs are shown from the side and from the extracellular space. Blue sticks: F703, red sticks: L704, purple sticks: F1374 and F1375, orange spheres: Cα of L695 and Y1365, teal spheres Cα of F582 and A1252.

**b.** The distance distribution between the Cα atoms of selected residue pairs (695/1365 and 582/1252) indicated that the transport pathway in the L704Y mutant of ABCG36 was comparable in width to that in the WT ABCG36. Therefore, the accelerated IAA translocation through the extracellular gate is most likely can be attributed to the high energy (FES) when IAA is in the binding pocket.

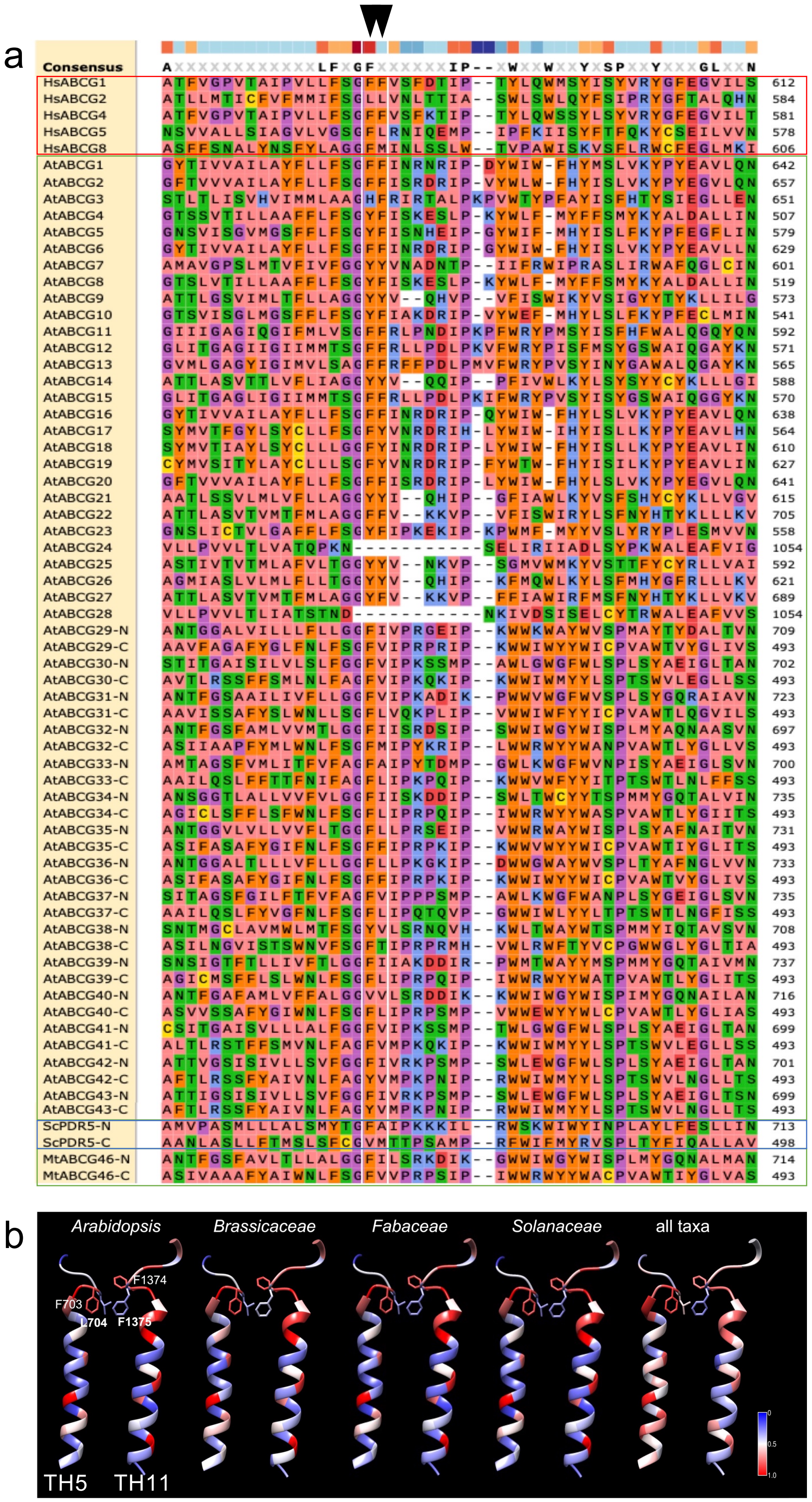


**Supplementary Figure 7: Conservation of essential residues in the extracellular gate of plant PDR-type ABCGs.**

**a.** Comparison of a proposed extracellular gate in HsABCG2 (Khunweeraphong et al., 2020; Huang et al., 2023; Ying et al., 2023) with homologous residues in Arabidopsis, human and selected other ABCGs. The position of two leucines, L554 and L555, forming the “di-leucine valve” in ABCG2 (Khunweeraphong et al., 2020), is boxed and indicated by a triangle. Human PDRS are boxedin red, yeasts ones in blue.

**b.** Conservation of indicated key residues of the extracellular gate in Arabidopsis. *Brassicaceae*, *Fabaceae* and *Solanaceae* families with all available sequences of plant full-size ABCG (PDR) sequences. Conservation was mapped on TH5 and TH11 and adjacent key residues corresponding to Arabidopsis F703/L704 and F1374/F1375 .

| | |
|---|---|
| L704R | FW: 5'-TTC CCC TTA GGG AGT CTA AAA CCA CCC AAA AGA AAT ACC AAC AAC A-3'<br>RV: 5'-TGT TGT TGG TAT TTC TTT TGG GTG GTT TTA GAC TCC CTA AGG GGA A-3' |
| L704D | FW: 5'-ATT TTC CCC TTA GGG AGA TCA AAA CCA CCC AAA AGA AAT ACC AAC AAC AAC-3'<br>RV: 5'-GTT GTT GTT GGT ATT TCT TTT GGG TGG TTT TGA TCT CCC TAA GGG GAA AAT-3' |
| L704S | FW: 5'-TTC CCC TTA GGG AGT GAA AAA CCA CCC AAA AGA AAT ACC AAC AAC A-3'<br>RV: 5'-TGT TGT TGG TAT TTC TTT TGG GTG GTT TTT CAC TCC CTA AGG GGA A-3' |
| L704C | FW: 5'-GAT TTT CCC CTT AGG GAG GCA AAA ACC ACC CAA AAG AAA TAC CAA CAA CAA CGT-3'<br>RV: 5'-ACG TTG TTG TTG GTA TTT CTT TTG GGT GGT TTT TGC CTC CCT AAG GGG AAA ATC-3' |
| L704A | FW: 5'-CCC CTT AGG GAG TGC AAA ACC ACC CAA AAG AAA TAC CAA C-3'<br>RV: 5'-GTT GGT ATT TCT TTT GGG TGG TTT TGC ACT CCC TAA GGG G-3' |
| L704I | FW: 5'-CCC CTT AGG GAG TAT AAA ACC ACC CAA AAG AAA TAC CA-3'<br>RV: 5'-TGG TAT TTC TTT TGG GTG GTT TTA TAC TCC CTA AGG GG-3' |
| L704Y | FW: 5'-GAT TTT CCC CTT AGG GAG ATA AAA ACC ACC CAA AAG AAA TAC CAA CAA CAA CGT-3'<br>RV: 5'-ACG TTG TTG TTG GTA TTT CTT TTG GGT GGT TTT TAT CTC CCT AAG GGG AAA ATC-3' |
| L704W | FW: 5'-GAT TTT CCC CTT AGG GAG CCA AAA ACC ACC CAA AAG AAA TAC CAA CAA CAA CGT-3'<br>RV: 5'-ACG TTG TTG TTG GTA TTT CTT TTG GGT GGT TTT TGG CTC CCT AAG GGG AAA ATC-3' |

**Table S1. Primer sequences for QuikChange site-directed mutagenesis**
